# A single small RNA shapes multiple symbiotic traits in rhizobia

**DOI:** 10.1101/2025.11.14.688489

**Authors:** Natalia I. García-Tomsig, Sabina K. Guedes-García, Marta Robledo, José I. Jiménez Zurdo

**Affiliations:** Structure, Dynamics and Function of Rhizobacterial Genomes (RhizoRNA Lab), Estación Experimental del Zaidín, Consejo Superior de Investigaciones Científicas (CSIC), Granada, Spain; Instituto de Biomedicina y Biotecnología de Cantabria, Universidad de Cantabria-CSIC, Cantabria, Santander, Spain

**Keywords:** host-microbe interactions, nitrogen fixation, nodulation, sRNA, NfeR1, cell cycle, bacteroid differentiation, *Sinorhizobium meliloti*

## Abstract

Bacterial small non-coding RNAs (sRNAs) remain understudied in the ecologically crucial nitrogen (N_2_)-fixing root-nodule *Rhizobium*-legume symbiosis. The only known rhizobial RNA regulator with broad symbiotic influence is the N-responsive *trans*-acting sRNA NfeR1, identified in the alfalfa symbiont *Sinorhizobium meliloti*. To pinpoint NfeR1 function, we profiled its RNA targets using MS2 affinity purification coupled with RNA sequencing (MAPS) in N stressed bacteria, a condition that drives nodulation. NfeR1 targets distinct regions of numerous mRNAs and sRNAs via three redundant anti-Shine-Dalgarno motifs, with silencing as major regulatory outcome. Target mRNAs span pathways differentially regulated throughout symbiosis, including N metabolism, motility, stress adaptation, and cell cycle control. Notably, NfeR1 modulates cell morphology and DNA replication by pervasive regulation of cell cycle mRNAs. It also silences *gdhA*, repressing glutamine dehydrogenase-dependent N assimilation and enhancing nodulation gene expression, further fine-tuned by a novel RNA feedback loop between NfeR1 and the dual-function sRNA SmelC549. Our findings position NfeR1 as a central hub within a structurally and functionally complex RNA network that coordinates N signaling and symbiotic performance in *S. meliloti*.

## Introduction

Regulation of gene expression by base-pairing small non-coding RNAs (sRNAs) provides bacteria with a ubiquitous mechanism for the swift and accurate fine-tuning of their transcriptional and translational outputs in response to environmental changes (Waters and Storz, 2009; Papenfort and Melamed, 2023). Specifically, sRNAs of the *trans*-acting class typically rely on short and discontinuous antisense interactions to the translation initiation region (TIR) of their target mRNAs to downregulate protein and/or mRNA abundance (Vogel and Wagner, 2007). In most cases, RNA chaperones such as Hfq or ProQ are required to facilitate this imperfect base pairing, which leads to regulation of large arrays of functionally related target mRNAs by a single sRNA (Quendera *et al*, 2020). *Trans*-sRNAs shape virtually all traits of bacterial physiology, notably those expressed during the chronic infection of a eukaryotic host (Chakravarty and Massé, 2019). The mutualistic root-nodule endosymbiosis between legume plants and soil dwelling diazotrophic α– and β–protobacteria, *i.e.* rhizobia, is a remarkable example of an intensively investigated beneficial host-microbe interaction because of its major impact in planet sustainability (Fowler *et al*, 2013). From a fundamental biology perspective, this symbiosis provides an invaluable experimental system for deciphering the subtleties of gene expression reprograming undergone by bacteria during plant infection (Jones *et al*, 2007; Haag *et al*, 2012). We have already anticipated the influence of some *trans*-sRNAs in the establishment of the symbiosis between the α–rhizobium *Sinorhizobium meliloti* and alfalfa (*Medicago sativa* L.) (Robledo *et al*, 2020; García-Tomsig *et al*, 2022, 2023; Robledo *et al*, 2017).

Nitrogen (N) scarcity is the primary abiotic cue in soil boosting symbiosis. Accordingly, the coordination of N assimilation and N_2_ fixation in nodules to support plant growth is a major metabolic adaptation experienced by rhizobia throughout the symbiotic interaction (Patriarca *et al*, 2002). The development of the N_2_-fixing root nodules relies on a precisely coordinated exchange of biotic signals between the symbiotic partners. Rhizospheric rhizobia first respond to root-exuded flavonoids to synthetize and secrete lipo-chitooligosaccharides (LCOs or Nod factors) that serve as elicitors of nodule organogenesis in a largely host-specific manner (Yu and Zhu, 2025; Jones et al, 2007; Haag et al, 2012). Upon root hair infection via the so-called infection threads and subsequent release into the cells of nodule primordia, rhizobia start a profound morphological differentiation to the form of N_2_-fixing competent bacteroids (Haag *et al*, 2012; Jones *et al*, 2007). Irreversible terminal bacteroid differentiation is a cell cycle-dependent process, which in members of the Inverted Repeat Lacking Clade (IRLC) of the Papilionoideae subfamily such as *M. sativa* L., is induced by nodule-specific cysteine-rich (NCR) peptides secreted by the plant (Van De Velde *et al*, 2010; Penterman *et al*, 2014; Haag *et al*, 2012; Jones *et al*, 2007). Mature bacteroids are accommodated intracellularly within symbiosomes that ensure the microaerobic environment required by nitrogenase to catalyze the reduction of atmospheric N_2_ to ammonia, which can then be incorporated into plant biomass (Jones *et al*, 2007; Patriarca *et al*, 2002; Haag *et al*, 2012). We have previously reported on an Hfq-independent N-responsive *S. meliloti trans*-sRNA termed NfeR1 (<u>N</u>odule <u>F</u>ormation <u>E</u>fficiency <u>R</u>NA) that is also highly expressed *in planta* throughout symbiosis with alfalfa (Robledo *et al*, 2017; García-Tomsig *et al*, 2023). Lack of NfeR1 delays nodulation kinetics, alters nodule organogenesis and impairs the overall symbiotic efficiency of *S. meliloti* on alfalfa roots. The *nfeR1* knockout mutant induces a large proportion of round-shaped and smaller nodules compared to the elongated, wild-type indeterminate mature alfalfa nodules. Remarkably, this represents the only reported genuine symbiotic phenotype linked to a sRNA loss-of-function in rhizobia, underscoring the potentially widespread regulatory role of NfeR1 in legume nodulation (Robledo *et al*, 2017, 2020).

Differential NfeR1 expression primarily relies on the LysR-type symbiotic regulator LsrB and the master regulator of the N stress response (NSR) NtrC, acting antagonistically as activator and repressor of transcription, respectively (García-Tomsig *et al*, 2023). Of note, the NtrBC two-component system and NfeR1 form a mixed protein-RNA double-negative feedback loop that strengthens both the NSR in free-living *S. meliloti* and the silencing of N assimilation by endosymbiotic bacteroids (García-Tomsig *et al*, 2023). NfeR1 hampers NtrB translation and promotes *ntrBC* decay upon a canonical interaction at the *ntrB* ribosome-binding site (RBS) *via* an ultraconserved anti-Shine-Dalgarno (aSD) motif (CCUCCUCCC) (García-Tomsig *et al*, 2023). This seed-pairing domain is present in each of the three hairpins in which NfeR1 and its α–proteobacterial orthologs, known as ‘CuKoo’ sRNAs, most likely fold (Reinkensmeier and Giegerich, 2015; del Val et al, 2012). Remarkably, only the concurrent mutation of all three aSD motifs disables NfeR1 for target mRNA regulation, indicating their functional redundancy (García-Tomsig *et al*, 2023; Robledo *et al*, 2017).

The antisense interaction at the RBS is highly effective in promoting post-transcriptional mRNA silencing (Vogel and Wagner, 2007; Waters and Storz, 2009). However, this mRNA element provides scarce sequence diversity and, therefore, supports a limited specificity for target regulation. As a result, *trans*-sRNAs employing ‘CuKoo’-like regulatory motifs have particularly large mRNA targetomes. This is the case of the sibling *S. meliloti* AbcR1 and AbcR2 *trans*-sRNAs, which use alike targeting aSD domains to interact with more than 200 metabolic mRNAs (García-Tomsig *et al*, 2022; Overlöper *et al*, 2014). As expected, computational predictions anticipate a similarly large NfeR1 regulon, with subsets of target mRNAs likely shared with AbcR1/2. In this study, we employed MS2 affinity purification combined with RNA sequencing (MAPS) to profile the RNA species that interact with NfeR1 *in vivo* upon bacterial growth in the N-limiting conditions prevailing at the onset of nodulation (Lalaouna *et al*, 2017). This genome-wide approach revealed arrays of mRNAs and sRNAs as specific NfeR1 partners. We further show that NfeR1-mediated regulation of some of these transcripts influences *S. meliloti* N metabolism, cell cycle progression and *nod* gene expression. Our data thus highlight a previously unrecognized, multifaceted role for this rhizobial sRNA in regulating key traits of legume nodulation.

## Results

### NfeR1 tagging and MAPS setup

To delve deeper into NfeR1 function we performed MAPS with a transcript tagged at the 5’-end with the MS2 aptamer (Fig. S1). Intriguingly, Northern blot probing of bacteria expressing MS2NfeR1 revealed accumulation of both wild type and tagged RNA species at similar levels. In this construct, NfeR1 was preceded by an *Xba*I recognition site, which is followed by the first TC nucleotides of *nfeR1*, yielding a methylation-sensitive GATC site (Hermann and Jeltsch, 2003). We reasoned that this site might either alter expression or promote MS2NfeR1 processing. Thus, we replaced a single or three alternative nucleotides within the *Xba*I site to generate MS2NfeR1_1 and MS2NfeR1_3, respectively, and probed RNA from bacteria expressing these variants. Only the triple nucleotide substitution completely prevented accumulation of the wild type NfeR1 transcript. Therefore, this version (hereafter referred to as NfeR1*) was used in the MAPS experiments.

To conduct MAPS, pSKiNfeR1 and pSKiNfeR1* were mobilized to strain Sm2020, which harbors *abcR1/2* and *nfeR1* deletions. Expression of wild type and tagged NfeR1 were induced for 15 min with isopropyl β-d-1-thiogalactopyranoside (IPTG) in transconjugants grown exponentially under N stress in MM (5.4 mM glutamate) or MM-NH_4_ (0.5 mM NH_4_Cl), and cell lysates were subjected to affinity chromatography (Fig. S2). Northern blot probing of RNA from the input and output chromatography fractions confirmed that the tagged sRNA was specifically retained in the columns (Fig. S2A). Mapping of the sequencing reads from the eluted RNA demonstrated efficient recovery of NfeR1* co-purified with known mRNA partners, *i.e*., *SMc03121*, *SMb20442* and *ntrB* (García-Tomsig *et al*, 2023; Robledo *et al*, 2017). Patterns of mapped reads on these mRNAs were similar in bacteria from MM and MM-NH_4_ cultures (Fig. S2B). RT-qPCR experiments further revealed that in the N-deficient MM, but not in rich TY broth, the first two mRNAs early identified as NfeR1 targets, *SMc03121* and *SMb20442*, were markedly upregulated in the SmΔ*nfeR1* mutant, indicating negative regulation by this sRNA (Fig. S2C). Together, these determinations validate the reliability of our experimental setup for the identification of the cellular transcripts interacting with NfeR1.

### Primary analysis of the NfeR1 targetome

To search for potential NfeR1 targets among the mRNAs retained in columns, we normalized by coverage and quantified sequencing reads across the full-length (FL) and the distinct regions of the mRNAs co-purified with NfeR1 and NfeR1*; TIR (spanning nucleotides -50 to +100 relative to the start codon), coding sequence (CDS), and virtual 3′-UTR (defined as 50 nt upstream and 30 downstream of the stop codon). We set a threshold of at least 100 mapped reads in any of these regions and imposed a minimum 2-fold difference (log_2_ FC>1) between NfeR1* and control NfeR1 samples to classify an mRNA as an NfeR1 target (Dataset S1). The mRNAs most likely captured because of complementarity to the MS2 aptamer sequence or enriched in regions overlapping other genomic features were not included in the candidates list. This analysis unveiled an exceptionally large set of 398 mRNAs likely targeted by NfeR1. Among these, 385 and 206 were identified upon bacterial growth in MM and MM-NH_4_, respectively, with 193 common to both conditions (Fig. 1A). Only 13 mRNAs, mostly encoding proteins of unknown function, were specifically retrieved under ammonia stress. Of all the target candidates, 67% encode proteins with predicted functions. Among these, 80% are likely involved in transport, metabolism, or transcriptional regulation (Fig. 1B). Of note, the genomic distribution of the targets, relative to the protein-coding gene content in each of the three *S. meliloti* replicons (chromosome, and symbiotic megaplasmids pSymA and pSymB), revealed a skew toward the chromosomal origin of these mRNAs (Fig. 1C).

**Figure 1.**
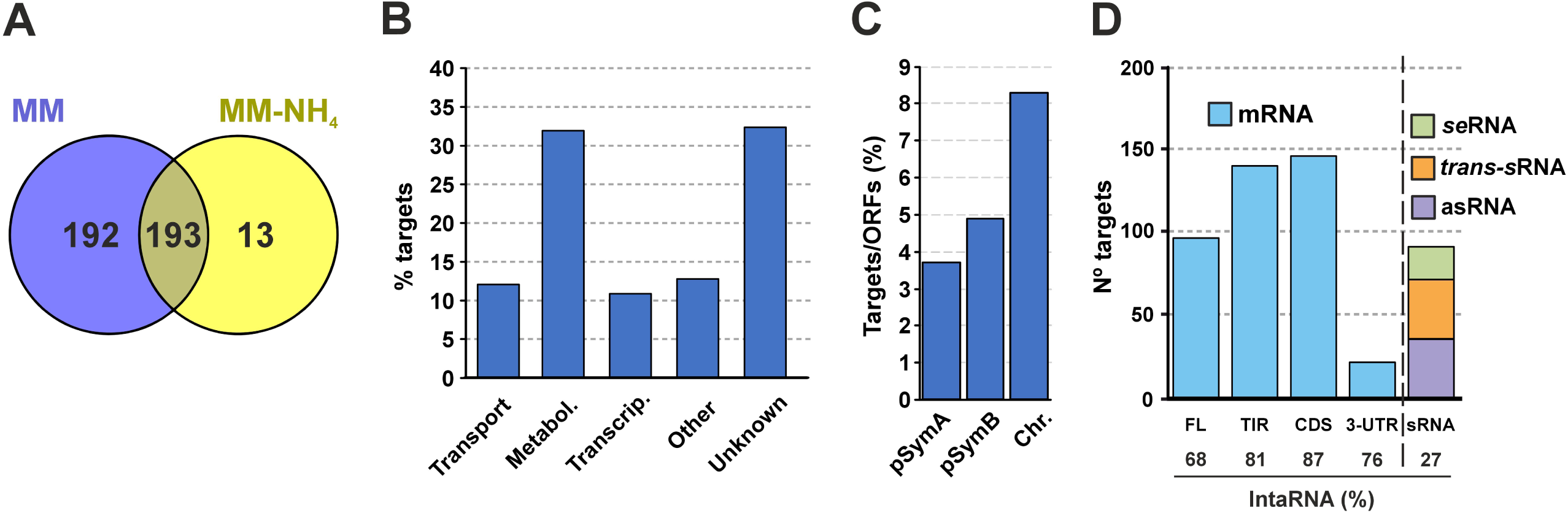
Overview of the NfeR1 RNA interactome uncovered by MAPS. (**A)** Venn diagram comparing the mRNA populations co-purified with NfeR1 in MM and MM-NH_4_. (**B)** Functional categories of the co-purified mRNAs. (**C)** Distribution of the target mRNA candidates relative to the total number of protein-coding genes in each of the three *S. meliloti* replicons: the pSymA and pSymB megaplasmids, and the chromosome. (**D)** Number of mRNAs and sRNAs co-purified with MS2NfeR1. The mRNAs are classified according to the region enriched in the procedure namely, the full-length transcript (FL), translation initiation region (TIR), coding sequence (CDS) and 3’-region (3’-UTR). Colours represent the different classes of sRNAs: sense sRNAs (seRNAs), antisense sRNAs (asRNAs) and trans-sRNAs. The *S. meliloti* genome was interrogated with IntaRNA to search for thermodynamically favourable antisense interactions (minimum 7-nt seed) of NfeR1 across the enriched mRNA region and full-length sRNAs. The percentages of positive predicted interactions are relative to the total number of target candidates within each category.

Read coverage biases in captured mRNAs often indicate the most likely interaction regions with the chromatography sRNA bait. Our analysis identified a large proportion of NfeR1 target mRNA candidates enriched in the TIR, CDS or both (FL mRNA), whereas enrichment solely in the virtual 3’-UTR was less frequent (Fig. 1D). Overall, IntaRNA predicted NfeR1 base pairing with approximately 87% of target mRNA candidates at the expected regions. These antisense interactions involve nucleotide seed sequences of seven or more residues and are thermodynamically favorable, with hybridization energies (*E*) below -8 kcal/mol.

Similarly, we profiled the population of sRNAs enriched during the procedure. This analysis identified 89 *S. meliloti* non-coding transcripts as NfeR1 targets, annotated as either sense mRNA-derived (seRNAs), antisense (asRNAs) or *trans*-acting sRNAs, with over 60% of those of chromosomal origin. IntaRNA predicted NfeR1 antisense interactions (*E* < -5 kcal/mol) for 16%, 37%, and 24% of seRNAs, asRNAs, and *trans*-sRNAs target candidates, respectively (Dataset S1). Nonetheless, base-pairing predictions may be missed for several of these transcripts due to incomplete sequence information, as their annotation is only based on determination of transcription start sites (Schlüter *et al*, 2013).

Our data thus reveal a broad NfeR1 RNA regulon that predominantly encompasses metabolic mRNAs and an unexpectedly large set of sRNAs. Further, predicted base-pairing interactions at multiple sites within the mRNA targets suggest alternative regulatory mechanisms distinct from the canonical translation inhibition via RBS occlusion.

### NfeR1 targets multiple pathways relevant to the symbiotic *S. meliloti* lifestyle

The nucleotide sequence of the NfeR1 aSD regulatory motifs closely resembles those of the sibling *trans*-sRNAs AbcR1 and AbcR2. However, scarcely 40% of the NfeR1 mRNA targets overlap with the previously reported combined AbcR1 and AbcR2 regulon (Fig. 2A). This set includes members of the *smo*, *agl*, *his* and *rha* operons, which encode ABC transporters (García-Tomsig *et al*, 2022). Like the *agl* operon, some other NfeR1 targets encode proteins related to the osmotic stress response, such as the transcription cleavage factor GreA, the pilin subunit PilA1, the transcriptional regulator FixK1 and the thiamin biosynthesis protein ThiC (Wei *et al*, 2004; Campbell *et al*, 2003; Vriezen *et al*, 2007; Rüberg *et al*, 2003). Among the 244 mRNAs uniquely targeted by NfeR1, we identified functional clusters related to N metabolism, motility and chemotaxis, and cell cycle pathways, which operate at different stages of the symbiotic interaction between *S. meliloti* and alfalfa (Table S1).

**Figure 2.**
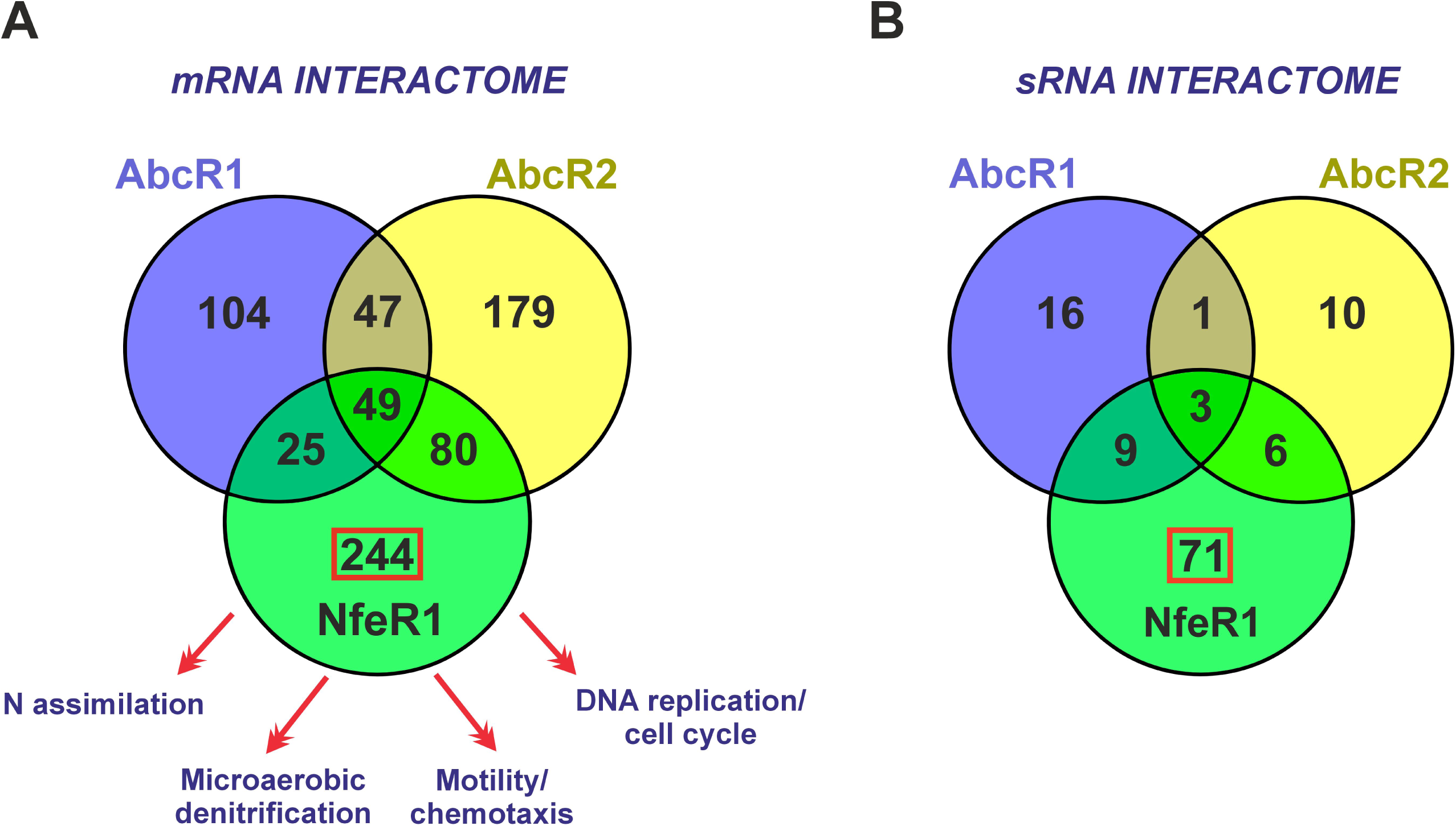
Comparison of the AbcR1, AbcR2 and NfeR1 RNA interactomes. (**A)** Venn diagram comparing mRNA targets. Symbiosis-related functional categories of specific NfeR1 target mRNAs are indicated. (**B)** Venn diagram corresponding to the sRNA targets.

In addition to *ntrB*, NfeR1 likely regulates other mRNAs involved in the N assimilation cascade active at the onset of nodulation, including the N sensors *glnB* and *glnE*, and the glutamate dehydrogenase *gdhA* (Yurgel et al, 2010; Dixon and Kahn, 2004; Mendoza et al, 1995). NfeR1 also targets transcripts encoding proteins involved in microaerobic denitrification, namely nitrous oxide reductase (*nosZ*), periplasmic nitrate reductase subunits (*napF* and *napE*), and the copper chaperone *nosL* (Torres *et al*, 2014; Pacheco *et al*, 2023). Motility and chemotaxis, which shape rhizobial behavior in the legume rhizosphere, also appear to be under NfeR1 control. Probable targets include mRNAs encoding regulatory and structural components of the flagellar machinery (e.g., *visN*, *rem*, *fliM*, *flgL*), and broad-range carboxylate chemoreceptors such as *mcpT* (Rotter *et al*, 2006; Baaziz *et al*, 2021; Sourjik *et al*, 2000). Furthermore, NfeR1-specific targets are enriched in transcripts related to DNA replication, repair, and cell division, suggesting a role in modulating cell cycle progression. Interestingly, we also observed a substantial enrichment of putative NfeR1 sRNA targets compared to those associated with the AbcR1/2 targetomes (Fig. 2B), identifying 29 asRNAs, 19 senseRNAs and 23 *trans*-sRNAs as NfeR1-specific targets, all of yet unknown function (Dataset S1). Together, these findings support a pleiotropic role for NfeR1 in fine-tuning *S. meliloti* physiology and symbiotic performance.

### Pervasive regulation of cell cycle mRNAs by NfeR1

A notable subset of putative NfeR1 targets includes mRNAs associated with the bacterial cell cycle (Table S1). Among these, we further investigated the influence of NfeR1 on the expression of *dnaA*, *divJ*, *smc*, *ftsK2*, *ftsY*, *ftsZ2* and *minC*. These mRNAs encode proteins likely operating at successive stages of the *S. meliloti* cell cycle progression: initiation of chromosomal replication (DnaA), control of replication timing (DivJ), chromosome organization (SMC), septum formation (FtsK2), membrane protein biogenesis (FtsY), and cell division (FtsZ2, MinC) (Pini et al, 2013; De Nisco et al, 2014; Cheng et al, 2007; Sibley et al, 2006; Gill and Salmond, 1987; Latch and Margolin, 1997; Yano and Niki, 2017; Veiga et al, 2023).

Analysis of sequencing read distributions on these mRNAs upon affinity chromatography revealed distinct enrichment patterns (Fig. 3A): 5’-biased for *dnaA*, 3’ enrichment for *ftsK2* and *divJ*, and uniform coverage along the entire *smc*, *ftsZ2*, *ftsY* and *minC* transcripts. Accordingly, IntaRNA predictions indicated favorable NfeR1 base pairing at either the TIR (*dnaA*) or disparate segments within the mRNA CDS (*ftsK2*, *divJ*, *smc*, *ftsZ2*, *ftsY* and *minC*) (Fig 3A, green arrows).

**Figure 3.**
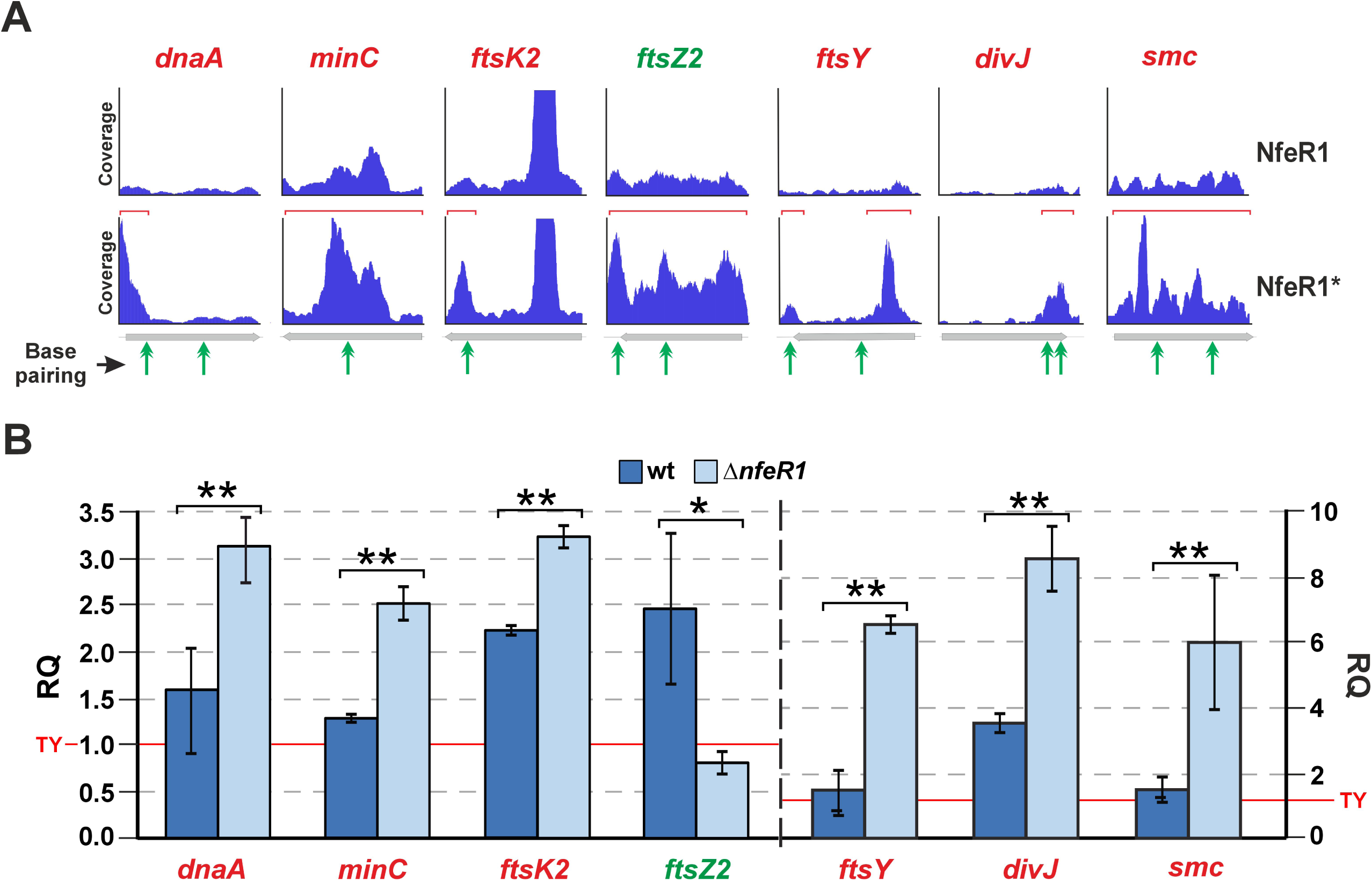
Regulation of cell cycle mRNAs by NfeR1. (A) Recovery profiles of the *dnaA*, *minC*, *ftsK2*, *ftsZ2*, *ftsY*, *divJ* and *smc* upon affinity chromatography with wild-type and tagged NfeR1 baits (IGV plots). Green arrowheads indicate the position of the predicted NfeR1 interaction site. (**B)** RT-qPCR analysis of mRNA abundance in the wild-type Sm2B3001 strain and the SmΔ*nfeR1* mutant. RNA was extracted from bacteria grown in TY to exponential phase, washed in PBS and then cultured in MM for 4 h. Relative Quantification (RQ) values were normalized to *SMc01852* as a constitutive control. The RQ values obtained from the MM cultures where related to the values obtained for the wild-type strain cultured in TY (red line). Bar graph values are the mean ± standard error of three replicates of two independent cultures (n = 6). Asterisks in the plots indicate significant differences according to ANOVA test, *p* < 0.01 (*) or *p* < 0.001 (**).

To assess NfeR1-mediated regulation of these mRNAs wild type and SmΔ*nfeR1* bacteria were first precultured in TY broth to log phase. N stress was then imposed to induce endogenous NfeR1 expression by suspending the cells in MM followed by a 4-h incubation. RT-qPCR analysis of RNA from these N-stressed bacteria revealed upregulation of the mRNA targets in the SmΔ*nfeR1* mutant compared to the wild-type strain, with the exception of *ftsZ2*, which was downregulated (Fig. 3B). As expected, no significant differences in levels of mRNA targets were detected between the two strains when cultured in TY, consistent with NfeR1 repression in nutrient-rich medium (Fig. S3). Furthermore, we also noted a significant induction of *ftsK2*, *ftsZ2* and *divJ* mRNAs in MM as compared to TY cultures, suggesting that expression of these genes is also N stress-dependent. These findings support that NfeR1 mediates broad post-transcriptional silencing of cell cycle-related mRNAs under N stress, while also suggesting a potential positive regulatory effect on *ftsZ2* via a distinct, yet uncharacterized mechanism.

### NfeR1 influences *S. meliloti* morphology and cell cycle progression under N stress

Misregulation of certain cell cycle-related NfeR1 targets is known to cause aberrant cell morphologies in *S. meliloti*, including branching, budding, or swelling (Latch and Margolin, 1997; Sadowsk et al, 2013; Sibley et al, 2006; Pini et al, 2015). To investigate potential NfeR1-dependent morphological phenotypes, we analysed TEM images of wild type and SmΔ*nfeR1* cells grown in MM (N stress) or TY broth to log phase (OD₆₀₀ 0.8) (Fig. S4). Approximately 2% of wild-type cells (n > 1000) exhibited abnormal features, most frequently Y-shaped, along with buds and non-polar branches.

These nuanced phenotypes were absent in bacteria grown in rich TY medium. In a comparable population, scarcely 1% of SmΔ*nfeR1* mutant cells displayed branching and budding morphologies upon growth in MM. Interestingly, IPTG-induced overexpression of NfeR1 from the pSKiNfeR1 plasmid triggered Y-shaped cell morphology in TY broth (0,5% cells; n = 500). In contrast, no morphological abnormalities were observed in the same strain without IPTG induction, nor in wild type cells grown in TY broth supplemented with IPTG, indicating that IPTG itself does not influence cell shape.

Branching events have been associated with various disruptions in *S. meliloti* cell cycle progression, including alterations in DNA replication (Pini *et al*, 2015; Sadowsk *et al*, 2013). Transcriptomics further reveal that several NfeR1-specific targets exhibit cell cycle-dependent expression patterns, such as *ftsK2*, *divJ*, and *minC*, or the motility and chemotaxis-related mRNAs *mcpT*, *rem*, *fliM*, *fliK*, and *flgL* (De Nisco *et al*, 2014). To further investigate the role of NfeR1 in this process, wild-type and SmΔ*nfeR1* strains were synchronized in G1 phase through N and carbon deprivation. Cells were then resuspended in MM to induce N stress, and harvested at 0, 2, 4, 6, and 16 hours. DNA content was assessed by fixing and staining cells with 4′,6-diamidino-2-phenylindole (DAPI) (Fig. 4). By 6 hours, wild-type cultures exhibited two distinct peaks corresponding to cells with one and two DNA copies (1C/2C). In contrast, the SmΔ*nfeR1* mutant displayed only a single 1C peak, indicating delayed DNA replication and segregation. Additionally, a substantial fraction of mutant cells showed faint DAPI staining, likely reflecting reduced viability. This interpretation aligns with technical difficulties encountered during harvesting of mutant bacteria, particularly as unstable pellets during washing, and suggests compromised viability during N and phosphate starvation in the absence of NfeR1.

**Figure 4.**
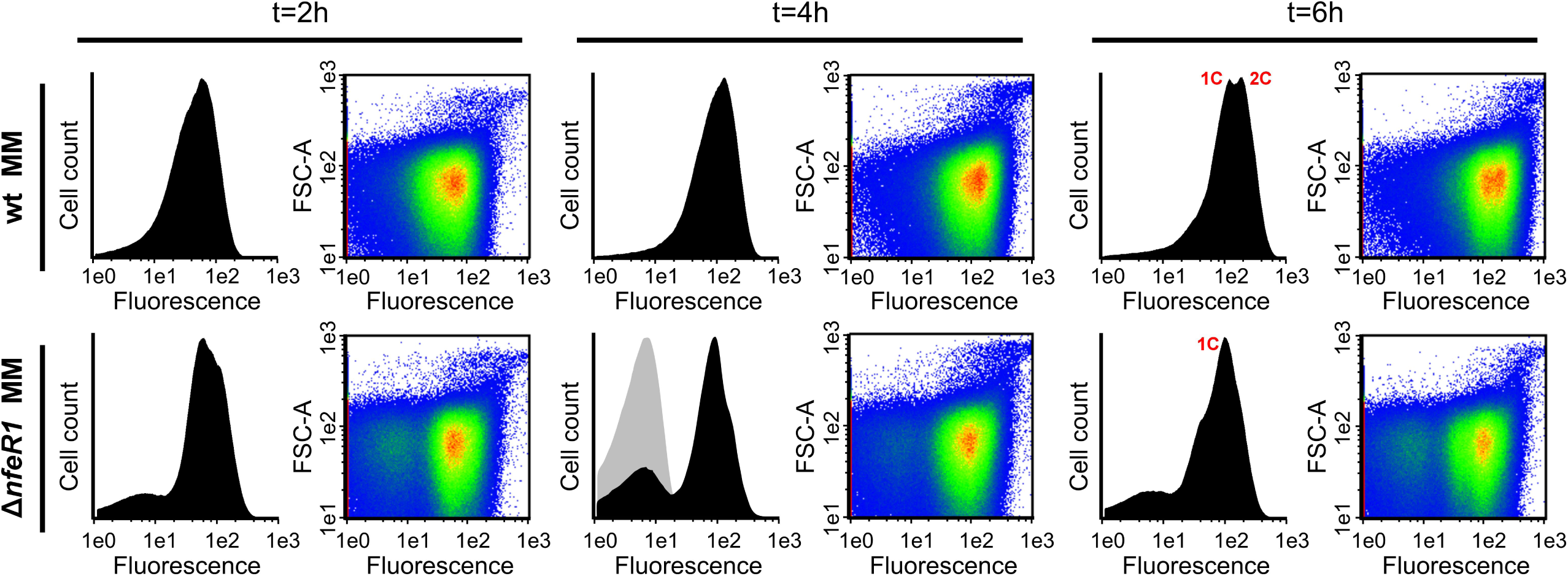
NfeR1 influences *S. meliloti* cell cycle progression. DNA content of wild-type Sm2B3001 and mutant SmΔ*nfeR1* bacteria measured by flow cytometry. Synchronized cultures were resuspended in MM, and cells were collected at the indicated time points. After DAPI staining, DNA content was quantified as fluorescence intensity. Left panels show DNA content (X-axis) versus cell count (Y-axis), with 1C and 2C indicating one and two genome equivalents, respectively. Right panels display density plots correlating DNA content with forward scatter (FSC).

To further confirm the impact of NfeR1 in cell cycle progression, the Sm2019Δ*nfeR1* mutant carrying plasmid pSKiNfeR1 was cultured in TY broth and sampled at 1, 2, 3, 4, and 16 hours following ectopic IPTG-induced expression of NfeR1 (Fig. S5). Distinct 1C and 2C DNA peaks were observed at 3 hours post-induction, whereas in non-induced cultures, the second peak emerged at 4 hours, suggesting that NfeR1 accelerates DNA replication. In contrast, a strain harboring plasmid pSKiNfeR1^abc^, which expresses the NfeR1 variant disabled for regulation, showed no differences in DNA content regardless of IPTG treatment, ruling out nonspecific effects of IPTG. Collectively, these findings highlight the significant influence of NfeR1 on *S. meliloti* cell cycle progression under N stress, likely through the promotion of DNA replication.

### NfeR1 underpins *nod* gene expression through *gdhA* silencing

Glutamate dehydrogenase (GDH) defines a minor N assimilation pathway that operates secondary to the predominant NtrBC-dependent glutamine synthetase (GS)-glutamate synthase (GOGAT) cascade (Dixon and Kahn, 2004; Arcondéguy et al, 2001). In *S. meliloti* 1021, GDH is encoded by the single-copy gene *gdhA*, located on the pSymA megaplasmid, yet considered part of the core genome of this species (Galibert *et al*, 2001). MAPS analysis identified *gdhA* mRNA as a target of NfeR1, showing enrichment along its entire length (Fig. 5A). IntaRNA predictions revealed that any of the three aSD motifs of NfeR1 could base pair with a 14-nt region within the *gdhA* CDS, each interaction exhibiting comparable hybridization energies (Fig. 5B). To further investigate NfeR1-mediated regulation of *gdhA*, we movilized plasmid p*P*_gdhA_*gdhA^FLAG^*, which expresses FLAG-tagged GdhA under its native promoter, into the SmΔ*nfeR1* strain carrying either pSKiNfeR1 or pSKiNfeR1^abc^. Upon IPTG induction in log-phase TY cultures, expression of wild-type NfeR1 led to a marked reduction in GdhA^FLAG^ levels, whereas the NfeR1^abc^ mutant had no effect (Fig. 5C). To assess GdhA regulation by the chromosomal NfeR1 copy, the Sm2019 strain expressing endogenous NfeR1 was transformed with pR*gdhA^FLAG^*, which drives constitutive production of GdhA^FLAG^. Western blot analysis of bacterial lysates revealed increased GdhA^FLAG^ levels in TY medium compared to MM, inversely correlating with known NfeR1 abundance under these conditions (Fig. 5D, left). RT-qPCR further confirmed elevated *gdhA* mRNA levels in SmΔ*nfeR1* relative to the wild-type strain, both in free-living cells subjected to N stress and in endosymbiotic nodule bacteria (Fig. 5D, right). These findings support that NfeR1 post-transcriptionally silences *gdhA* via base pairing of its redundant aSD motifs to the mRNA CDS.

**Figure 5.**
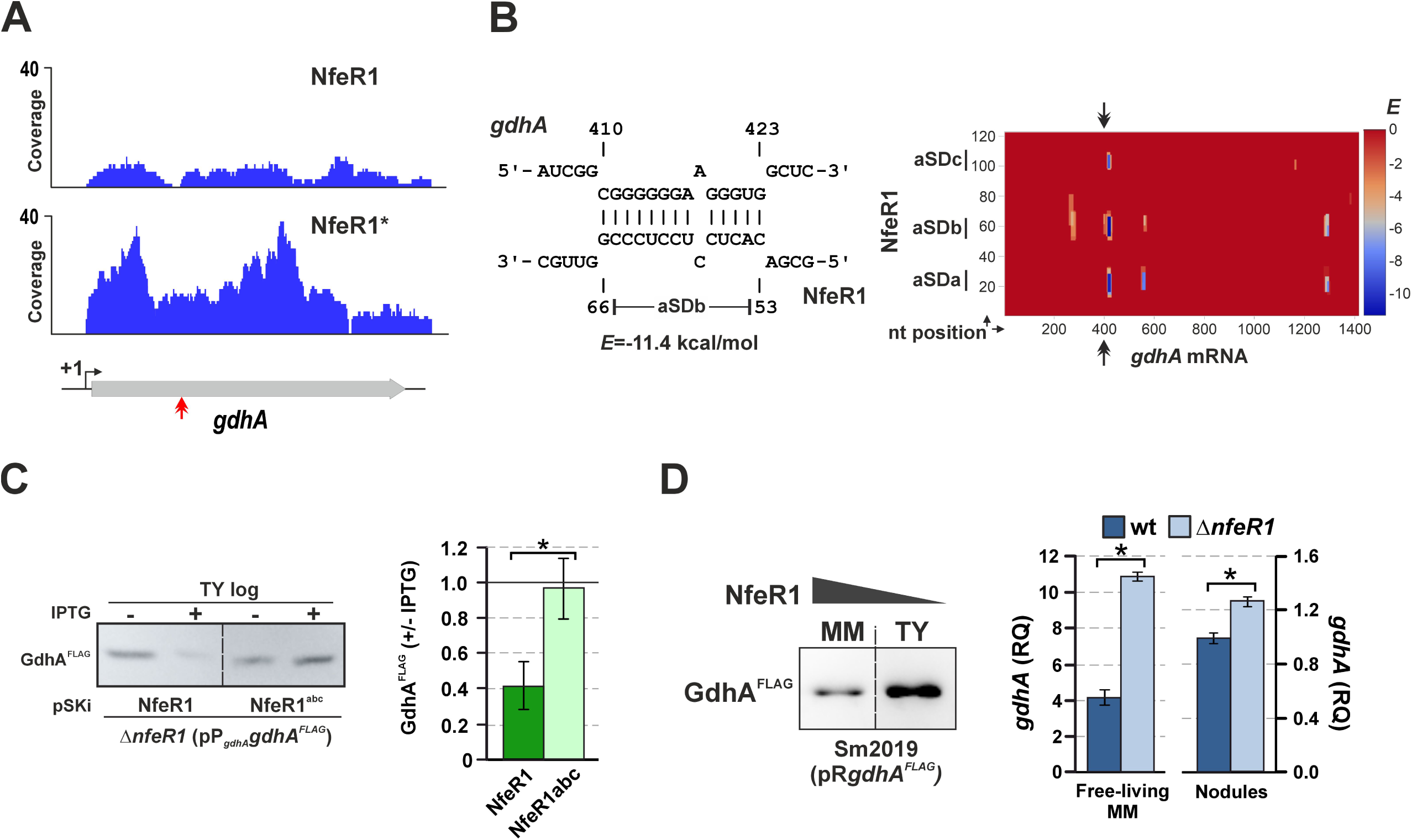
NfeR1 silences *gdhA*. (A) Recovery profiles of the *gdhA* mRNA following affinity chromatography with wild-type and tagged NfeR1 baits (IGV plots). The transcription start site (TSS) of *gdhA* is indicated (+1). The red double arrowhead marks the predicted NfeR1 interaction site. (B) IntaRNA-predicted base-pairing between NfeR1 and *gdhA* mRNA. Shown is the interaction with the aSDb motif (nt 53-66). Nucleotide positions in the mRNA are relative to the TSS. IntaRNA predicted similar *E* for hybridization with aSDa,c motifs (right graph). (**C)** NfeR1 downregulates GdhA. Western blot analysis of GdhA^FLAG^ produced from pP*_gdhA_gdhA^FLAG^* in Sm2019Δ*nfeR1* following IPTG-induced expression of native NfeR1 or the motif-mutated variant NfeR1abc from pSKi-derived plasmids. In p*P_gdhA_gdhA^FLAG^*, *gdhA^FLAG^* is transcribed from its native promoter (*P_gdhA_*). Gel lanes were loaded with equal protein amounts (OD_600_ equivalent to 0.2). GdhA^FLAG^ accumulation was quantified from band intensities 16 h after IPTG addition in three independent TY cultures of three independent double transconjugants. Samples were loaded twice as technical replicates (n = 6). Bar graph values are the mean ± standard error of the +/- IPTG ratios. Significant differences (*) were identified using ANOVA (*p*<0.05). (**D)** Regulation of *gdhA* by the endogenous NfeR1. Left: Western blot analysis of GdhA^FLAG^ produced from pR*gdhA^FLAG^* in log-phase Sm2019 MM and TY cultures. In pR*gdhA^FLAG^*, *gdhA^FLAG^*is transcribed from a constitutive promoter (*P_syn_*). Endogenous NfeR1 accumulation under these conditions is indicated above. Gel loading and replicates as in panel “C”. Right: RT-qPCR of *gdhA* in wild-type Sm2B3001 and SmΔ*nfeR1* strains. RNA derived from MM cultures or bacteria isolated from mature alfalfa nodules. Relative Quantification (RQ) values were normalized to *SMc01852* as a constitutive control. Bar graph values are means ± standard error from three replicates of two independent cultures (n = 6). Significant differences (*) were identified using ANOVA (*p*<0.001).

Ectopic (over)expression of heterologous *gdh* genes in rhizobia lacking assimilatory GDH activity interferes with *nod* gene expression and impairs symbiotic N_2_ fixation (Osburne and Signer, 1980; Mendoza et al, 1995). Thus, we next examined the impact of GdhA and NfeR1 on *nodA* mRNA accumulation (Fig. 6). As expected, addition of the alfalfa flavonoid luteolin to log-phase MM cultures of wild-type Sm1021 induced a 4-fold increase in *nodA* expression, while *gdhA* levels remained unchanged (Fig. 6, left). Notably, lack of NfeR1 led to reduced *nodA* induction upon luteolin treatment, an effect recapitulated by constitutive overexpression of *gdhA* in the wild-type background (Fig. 6, right). Importantly, deletion of *gdhA* in SmΔ*nfeR1* restored *nodA* expression to near wild-type levels (Fig. 6, left). These results indicate that full expression of *S. meliloti nod* genes requires NfeR1-mediated silencing of *gdhA*.

**Figure 6.**
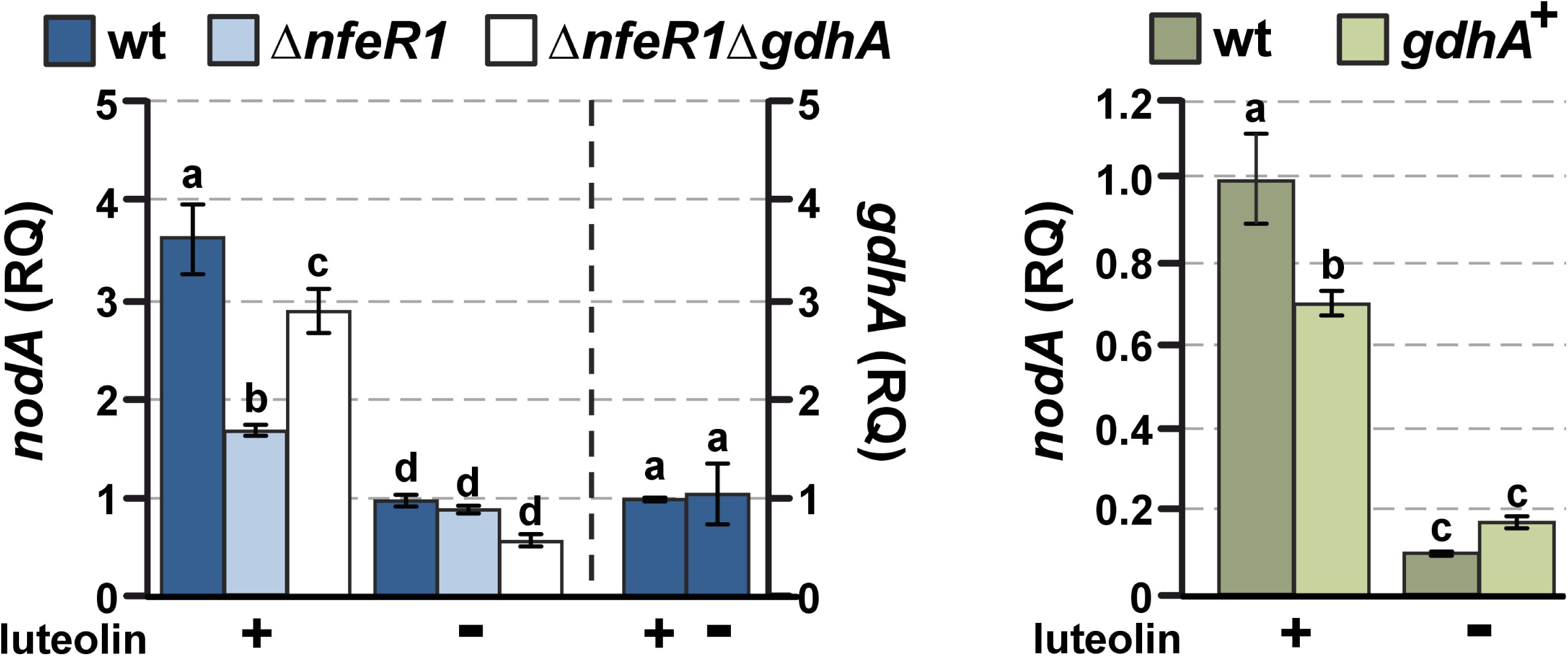
NfeR1-mediated *gdhA* silencing strengthens luteolin-induced *nodA* expression. RT-qPCR analysis of *nodA* mRNA levels in *S. meliloti* strains cultured in MM-NH_4_. Luteolin diluted in ethanol was added to log-phase cultures to induce *nod* gene expression. RNA was extracted 16 h after luteolin addition (+). As a control, ethanol alone was added to mock-treated cultures (-). Left: *nodA* mRNA abundance in wild-type Sm2B3001 bacteria, and SmΔ*nfeR1* and SmΔ*nfeR1*Δ*gdhA* derivatives. The *gdhA* mRNA levels in Sm2B3001 with or without luteolin were determined as a control. Right: *nodA* mRNA levels in Sm2B3001 carrying plasmid pSK*gdhA^FLAG^* for IPTG-induced overexpression of *gdhA.* IPTG was added 2 h prior luteolin or ethanol addition. Relative Quantification (RQ) values were normalized to *SMc01852* as a constitutive control. Bar graphs values are means ± standard error from three replicates of two independent cultures (n = 6). Different letters above bars indicate statistically significant differences between groups (one-way ANOVA, *p* < 0.05, followed by post hoc Tukey’s test).

### Mutual regulation between NfeR1 and the SmelC549 sRNA

Among the extensive set of NfeR1-specific sRNA targets, we focused on the 258 nt-long SmelC549 (SMc06721) for two main reasons: (i) it was recently catalogued as a dual-function sRNA that internally encodes the 25-aa small peptide SEPr6, and (ii) IntaRNA predictions indicated potential base pairing with nodulation-related mRNAs (Schlüter *et al*, 2010; Hadjeras *et al*, 2023). The 5′ region of SmelC549 (nt 1-189) shares sequence homology with the rhizobial sRNA Atu_C9 (RF02503), originally described in *Agrobacterium tumefacien*s, and is syntenic with the divergently transcribed *metF* gene (Fig. 7A), which encodes 5,10-methylenetetrahydrofolate reductase (Wilms *et al*, 2012). Notably, the SEPr6 coding sequence is highly conserved across several rhizobial species, including *S. medicae*, *S. fredii*, *Rhizobium leguminosarum*, *R. tropici*, and *R. etli*, whereas an alternative stop codon was detected in the *Brucella* homolog (Fig. S6A). IntaRNA analysis predicted energetically favourable base pairing (*E* < –8 kcal/mol; seed length 7 nt) between the RBS of *SEPr6* and the three aSD motifs of NfeR1 (Fig. 7B). This putative interaction is also conserved among homologous NfeR1-SmelC549 regulatory pairs in rhizobia, particularly in *S. medicae*, *S. fredii* and *R. tropici* (*E* **<** –10 kcal/mol; seed length 7 nt) (Fig. S6B).

**Figure 7.**
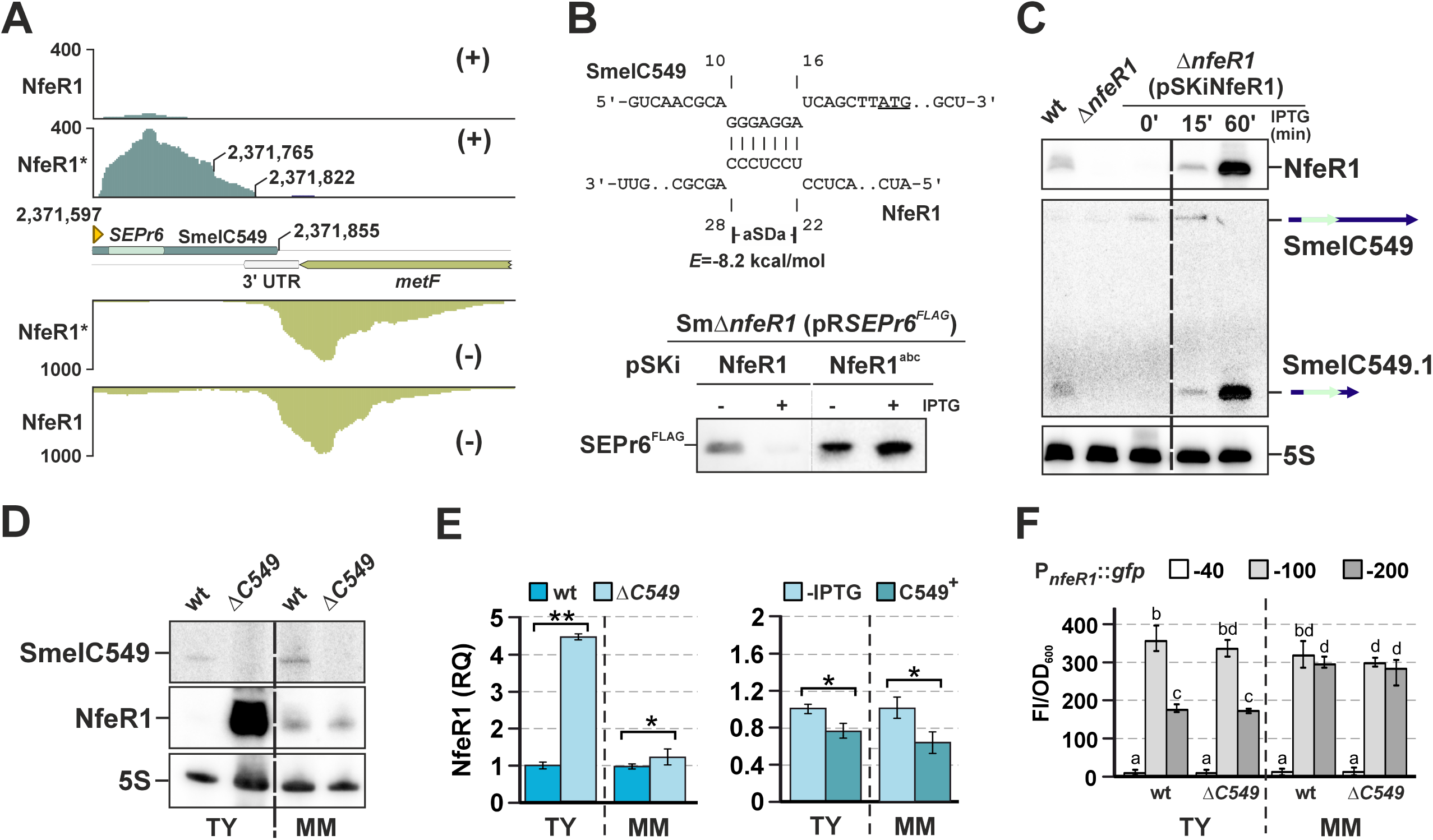
Mutual regulation between NfeR1 and SmelC549. **(A)** Recovery of SmelC549 and *metF* following affinity chromatography with wild-type and tagged NfeR1 baits. Gene coordinates refer to chromosomal positions. (**B)** Post-transcriptional regulation of SEPr6 by NfeR1. IntaRNA predicted base-pairing between NfeR1 aSDa and SmelC549. The SEPr6 start codon is underlined. Similar hybridization energies (E) were predicted for interactions with aSDb,c motifs. Bottom: Western blot analysis of SEPr6^FLAG^ produced constitutively from pR*SEPr6^FLAG^* in Sm2019Δ*nfeR1* after 16 h of IPTG-induced expression of NfeR1 or its mutant variant NfeR1abc from pSKi derivatives. Gel lanes were loaded with equal protein amounts (OD_600_ equivalent to 0.2). Three independent cultures from three independent double transconjugants were loaded twice into different gels as technical replicates (n = 6). (**C)** Northern blot analysis of NfeR1-dependent SmelC549 expression. Total RNA was extracted from MM cultures of Sm2B3001, SmΔ*nfeR1* and Sm2019Δ*nfeR1*, the latter carrying plasmid pSKiNfeR1 for IPTG-induced expression of NfeR1 at the indicated time points. (**D)** Northern blot analysis of SmelC549-dependent NfeR1 expression. Total RNA was extracted from Sm2B3001 and SmΔ*C549* cultured in TY and MM. The 5S rRNA was probed as loading control. (**E)** RT-qPCR analysis of NfeR1 in Sm2B3001 and SmΔ*C549* (left), and Sm2019Δ*C549* carrying plasmid pSKiC549 (right) cultured in TY and MM. Relative Quantification (RQ) values were normalized to *SMc01852* as a constitutive control. Bar graphs values are means ± standard deviation (n = 6). Asterisks in the plots indicate significant differences according to ANOVA: *p* < 0.05 (*) or *p*<0.001 (**). (**F)** Transcriptional regulation of NfeR1 assessed by promoter-*eGFP* fusions. Fluorescence derived from full-length (P*_nfeR1-_*_213_) and trimmed versions of P*_nfeR1_*(P*_nfeR1-_*_100_, P*_nfeR1-_*_40_) was measured in Sm2B3001 and SmΔ*C549* grown in TY or MM (n = 9). Different letters above bars indicate statistically significant differences between groups (one-way ANOVA, *p* < 0.05, followed by post hoc Tukey’s test).

To evaluate the influence of NfeR1 on SEPr6 translation, we performed Western blot analysis on total protein extracts from Sm2019Δ*nfeR1* cells co-transformed with pRP*_syn_SEPr6^FLAG^* (constitutively expressing SEPr6^FLAG^) and pSKiNfeR1 or pSKiNfeR1^abc^. The results showed that NfeR1 represses SEPr6 translation, whereas overexpression of NfeR1^abc^ did not affect protein accumulation (Fig. 7B). In parallel, Northern blot analyses revealed the accumulation of the full-length SmelC549 along with a shorter isoform (SmelC549.1), which was absent in the SmΔ*nfeR1* mutant, during log-phase growth of wild-type bacteria in MM (Fig. 7C). Furthermore, endogenous SmelC549.1 accumulation mirrored that of NfeR1 upon IPTG-induced expression from pSKiNfeR1, suggesting that NfeR1 may promote processing of the full-length SmelC549 transcript to this shorter more stable RNA species.

Strikingly, Northern blot analysis revealed that lack of SmelC549 led to increased accumulation of NfeR1 in rich TY broth, whereas this effect was not observed in MM cultures (Fig. 7D). RT-qPCR further confirmed this finding and additionally showed a slight decrease in NfeR1 levels upon IPTG-induced overexpression of SmelC549 from plasmid pSKiSmelC549 in the SmΔ*C549* background (Fig. 7E). To determine whether this regulation occurred at the transcriptional level, we analysed GFP reporter fusions of three NfeR1 promoter variants, P*_-40_*, P*_-100_*and P*_-200_*, containing predicted binding sites for RpoD, LsrB and NtrC/LsrB, respectively (Fig. 7F). As expected, P*_-40_::gfp* showed basal fluorescence, while LsrB-dependent activation of P*_-100_::gfp* was observed in both TY and MM media. Furthermore, NtrC-mediated repression of P*_-200_::gfp* was specifically detected in TY-grown cells as expected (García-Tomsig *et al*, 2023). Remarkably, the fluorescence levels of all promoter fusions remained unchanged in the SmΔ*C549* mutant, indicating that SmelC549 does not affect NfeR1 transcription. These findings thus suggest that SmelC549 regulates NfeR1 post-transcriptionally likely functioning as an RNA sponge. Taken together, the data support a cross-regulatory interaction between NfeR1 and SmelC549, revealing a novel and conserved RNA-based regulatory loop in rhizobia.

### SmelC549 promotes NodD2 accumulation under N stress

To begin uncovering the function of SmelC549 as base-pairing *trans*-sRNA, we used its full-length sequence to interrogate the *S. meliloti* genome with IntaRNA. This analysis delivered a broad set of putative targets, including *nodD1*, *nodD2*, and *nodD3* mRNAs (*E* < –10 kcal/mol; seed length: 7 nt). NodD proteins are LysR-type transcriptional regulators that respond to plant-derived flavonoids to activate *nod* gene expression in rhizobia (Ampe et al, 2003; Honma and Ausubel, 1987; Jones et al, 2007). Notably, an exceptionally long 141-nt interaction region was predicted within the *nodD2* mRNA (E < –25 kcal/mol), a feature not conserved in *nodD1* or *nodD3* (Fig. 8A and B).

**Figure 8.**
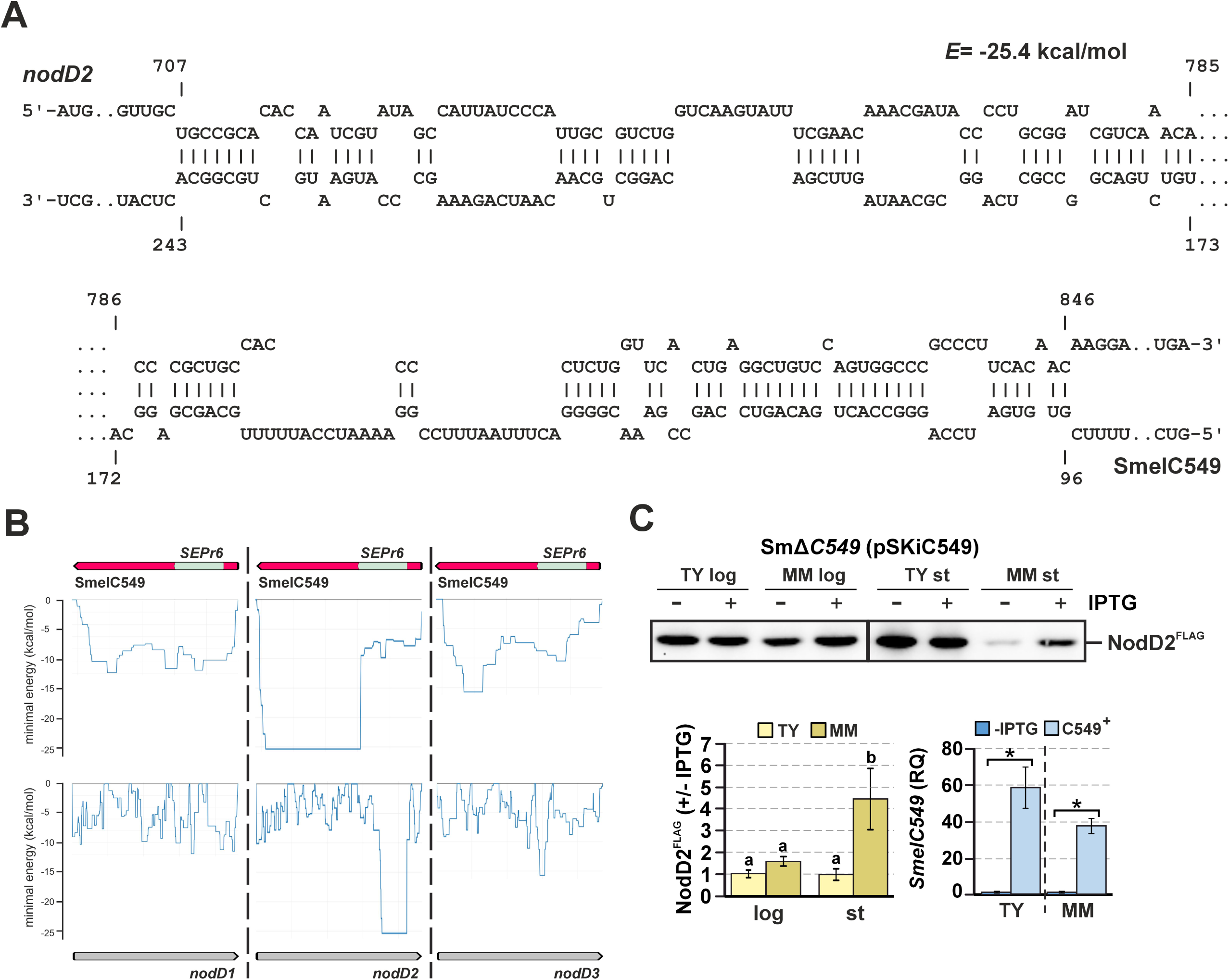
Post-transcriptional regulation of *nodD2* by SmelC549. (**A)** IntaRNA predicted base-pairing between SmelC549 and the *nodD2* mRNA. Nucleotide positions are indicated relative to the *nodD2* mRNA start codon. (**B)** Comparative IntaRNA predictions of SmelC549 interactions with *nodD1*, *nodD2* and *nodD3* mRNAs. (**C)** Upper panel: Western blot analysis of SEPr6^FLAG^ constitutively produced in Sm2019Δ*C549* after 4 h (log-phase cultures; log) or 16 h (stationary-phase cultures; st) of IPTG-induced SmelC549 transcription from pSKiC549. Gel lanes were loaded with equal protein amounts (OD_600_ equivalent to 0.2). Samples were loaded twice into separate gels as technical replicates (n = 6). Bottom left panel: bar graph showing mean ± standard error of the +/- IPTG ratios of band intensities. Different letters above bars indicate statistically significant differences between groups (one-way ANOVA, *p* < 0.05, followed by post hoc Tukey’s test). Bottom right panel: SmelC549 abundance in Sm2019Δ*C549* carrying plasmid pSKiC549 cultured in TY or MM. RNA was extracted 16 h after IPTG addition. Relative Quantification (RQ) values were normalized to *SMc01852* as a constitutive control. Bar graph values are mean ± standard error from three replicates of two independent cultures (n = 6). Asterisks (*) in the plots indicate significant differences inferred from ANOVA, *p* < 0.05.

To assess the functional relevance of this interaction, we performed Western blot analysis on Sm2019Δ*C549* cells co-transformed with pSKiSmelC549 and pR-*nodD2^FLAG^*, which constitutively expresses FLAG-tagged NodD2. After IPTG-induced expression (16 h), SmelC549 promoted NodD2 accumulation in MM, but not in TY broth (Fig. 8C). We also observed downregulation of NodD2^FLAG^ in stationary-phase MM cultures, potentially reflecting silencing of SmelC549 or the involvement of other regulatory factors affecting *nodD2* expression. Given that the processed SmelC549.1 species accumulates specifically in MM when NfeR1 is induced, the post-transcriptional regulation of *nodD2* likely involves this isoform. These findings uncover a previously unrecognized RNA-based regulatory layer within the complex *nod*-gene network of *S. meliloti*.

## Discussion

Although transcriptional regulation of symbiotic genes in rhizobia is well understood, post-transcriptional control remains largely unexplored. Here, we provide a comprehensive target-centric perspective of the network shaped by the conserved rhizobial N-responsive *trans*-sRNA NfeR1 in *S. meliloti*. Through base pairing at different regions, NfeR1 targets numerous mRNAs and sRNAs, fine-tuning a diversity of pathways active during the symbiotic transition (Fig. 9). By modulating the NtrBC signaling cascade, NfeR1 integrates N status to the requirements of symbiosis, underscoring its pleiotropic role in symbiotic performance. These findings provide the first evidence that a single *trans*-acting sRNA can exert broad regulatory control over diverse genuine symbiotic traits.

**Figure 9.**
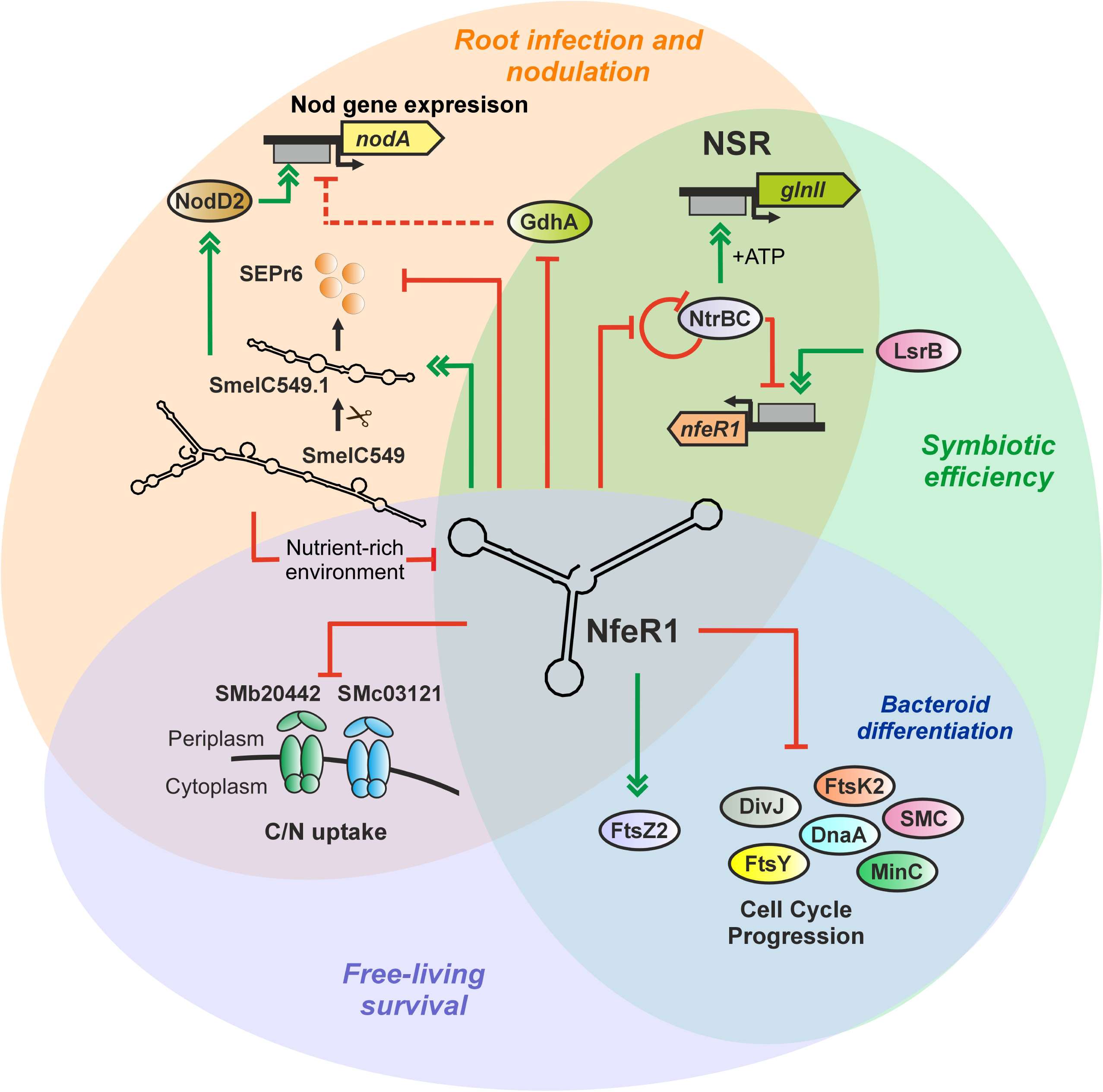
Multifaceted regulation of legume nodulation pathways by the *S. meliloti* sRNA NfeR1. Solid and dashed lines indicate direct and indirect regulation, respectively. Truncated red lines stand for negative regulation, whereas the green double arrowheads indicate positive regulation.

### MAPS-derived insights into the activity mechanism and symbiotic function of NfeR1

To delineate the function of NfeR1 in a symbiotic framework, we conducted a MAPS-based genome-wide profiling of its RNA interaction partners upon *S. meliloti* growth under N stress, the environmental condition that triggers legume nodulation (Patriarca *et al*, 2002). This approach revealed that NfeR1 likely targets approximately 7% of the protein-coding genes in the *S. meliloti* genome, primarily through the three redundant aSD regulatory motifs. Most of these mRNAs encode proteins involved in nutrient uptake and metabolism, with partial overlap observed with the previously characterized AbcR1/2 regulon. This was an expected finding, as all three sRNAs use aSD motifs with similar nucleotide sequences for target recognition. However, the primary analysis of the NfeR1 mRNA interactome revealed two major differences when compared to those of AbcR1 and AbcR2: (i) NfeR1 targets are more frequently located on the *S. meliloti* chromosome, suggesting a broader role for this sRNA in the evolutionary tuning of core functions to symbiotic conditions; and (ii) although MAPS profiles indicated a predominant canonical targeting at the mRNA TIR, interactions were equally predicted within CDSs. Indeed, the number of NfeR1 targets enriched in their coding regions far outnumbered that of AbcR1/2-interacting mRNAs with similar read distribution patterns after MAPS. Despite this diversity of interaction signatures, our data evidenced that mRNA silencing is the prevalent outcome of NfeR1-mediated regulation. The notable exception among the validated mRNA targets was *ftsZ2*, which is positively regulated by NfeR1 though predicted interactions deep within the CDS and the 3’-UTR. sRNA base-pairing outside the TIR, have been reported to differently influence both mRNA stability and translation efficiency (Overlöper et al, 2014; Papenfort and Vogel, 2009; Fröhlich et al, 2012; Rice and Vanderpool, 2011). Our dataset therefore represents a valuable new resource for exploring the mechanistic plasticity of sRNA-mediated regulation in rhizobia.

Comparison of the NfeR1 and AbcR1/2 regulons further revealed that NfeR1 specifically targets mRNAs involved in a wide range of cellular pathways including chemotaxis, flagella assembly, osmotic stress response, N metabolism, and cell cycle control. Chemoreceptors and flagella promote swarming and/or swimming motility, facilitating rhizobial colonization of the rhizosphere, root surfaces, and even nodule tissues (De Nisco *et al*, 2014; Sourjik *et al*, 2000; Amaya-Gómez *et al*, 2015; Rotter *et al*, 2006; Baaziz *et al*, 2021). However, *S. meliloti* strains lacking NfeR1 did not exhibit defects in motility or alfalfa root colonization, whereas AbcR1 has been shown to enhance colonization efficiency (Robledo *et al*, 2017; García-Tomsig *et al*, 2023). This phenotype further suggests that NfeR1 regulates unknown aspects of *S. meliloti* physiology that are distinct from those controlled by AbcR1/2.

We have also shown that NfeR1 contributes to salt/osmotic stress adaptation in *S. meliloti* (Robledo *et al*, 2017). Osmotolerance and the capacity to establish long-term intracellular residence within eukaryotic host cells are shared phenotypic traits of both mammalian pathogens and N_2_-fixing legume endosymbionts (Mahmoud et al, 2017; Poole and Ledermann, 2022; Hawkins and Oresnik, 2022). Notably, the expression of several NfeR1-interacting mRNAs such as those involved in the accumulation of osmoprotectants (e.g., trehalose), is hypersaline-dependent and contributes to both salt tolerance and effective symbiosis (Vriezen *et al*, 2007; Wei *et al*, 2004; Campbell *et al*, 2003; Rüberg *et al*, 2003; Ledermann *et al*, 2021). MAPS analysis further positioned NfeR1 as regulator of multiple N metabolism pathways, including N uptake, denitrification, N_2_ fixation, and aerobic N assimilation. In nodules, the loss of fixed N, either through uptake and reassimilation by bacteroids or through nitrate and nitrite reduction via upregulated microaerobic denitrification, can hinder plant growth (Patriarca et al, 2002; Haag et al, 2012; Jones et al, 2007; Dixon and Kahn, 2004). Moreover, nitric oxide, an intermediate in this process, is toxic and directly inhibits nitrogenase activity (Torres *et al*, 2014; Pacheco *et al*, 2023). Therefore, the widespread silencing of this set of mRNAs by NfeR1 in mature bacteroids may enhance overall symbiotic efficiency.

### NfeR1 as a regulatory node in adapting cell cycle progression to N stress and symbiosis

Bacteria dynamically modulate their cell cycle to match growth pace with environmental demands. This housekeeping process is tightly regulated by diverse, and often redundant, transcriptional, post-transcriptional, and post-translational mechanisms that remain poorly understood in most species (De Nisco et al, 2014; Macek et al, 2019; Meunier & Govers, 2025; Fröhlich and Velasco Gomariz, 2021). Our data show that NfeR1 broadly influences *S. meliloti* cell cycle progression by post-transcriptionally regulating multiple mRNAs predicted to be involved at various stages of this process. While EcpR1 and GspR, two other *S. meliloti* sRNAs, also target cell cycle regulators mRNAs such as *ctrA*, *dnaA*, and *gcrA*, they do not respond to N stress or affect nodulation and symbiosis, unlike NfeR1 (Robledo *et al*, 2015, 2018). These findings emphasize the central role of bacterial sRNAs in integrating diverse regulatory signals to shape core cellular processes and phenotypic outcomes.

Adaptation of rhizobial growth to N stress is primarily coordinated by the NtrY/NtrX two-component system, which transcriptionally regulates genes involved in both N metabolism and cell cycle control, including *ctrA*, *gcrA*, *dnaA* and *ftsZ1,* which is the *S. meliloti* homolog of the *Escherichia coli* and *A. tumefaciens ftsZ* (Xing et al, 2022; Margolin and Long, 1994). Notably, *ntrX* knockout and altered expression of certain NfeR1 target mRNAs, such as *dnaA*, *minC* and *ftsZ2* are associated with prevalent morphological alterations in the mutant populations, including elongation, budding, and Y- or V-shaped cells (Xing *et al*, 2022). Interestingly, we observed that N stress alone induces similar phenotypes in a minority of wild-type bacteria, suggesting that these morphological changes are likely a transient feature of the adaptive response. The absence of NfeR1 slightly altered this pattern under N stress, while its overexpression disrupted the homogeneity of the bacillary population, inducing stress-related morphologies even under mild culture conditions. Tracking DNA content in these loss- and gain-of-function mutants by flow cytometry evidenced that NfeR1 has a major impact on coordinating genome duplication with growth by promoting DNA replication. Besides metabolic reprogramming, NfeR1-driven modulation of cell cycle progression thus play an important role in helping *S. meliloti* withstand N stress. This regulatory mechanism provides a compelling additional explanation for the major N stress-related phenotypes observed in free-living NfeR1 mutant bacteria, specifically their weakened survival and adaptability.

Several lines of evidence indicate that modulation of the *S. meliloti* cell cycle is also critically important for successful plant infection and bacteroid differentiation (De Nisco et al, 2014; Xue and Biondi, 2019). Following root hair infection, rhizobia colonize the infection thread lumen through phases of rapid division and coordinated movement until their release into the cells of the nodule primordia. At this stage, cell division arrests and genome endoreduplication begins, leading to the formation of Y-shaped, oversized polyploid bacteroids that are competent for N_2_ fixation (Jones et al, 2007; Yu and Zhu, 2025; Haag et al, 2012). Notably, we reported that the small, round-shaped nodules induced by the *S. meliloti nfeR1* knockout mutant on alfalfa roots are filled with endosymbiotic bacteria. However, plant leghemoglobin, a marker for N_2_ fixation, is absent in most of these nodules or exhibits significantly delayed expression compared to those formed by the wild-type strain, indicating defects in bacteroid maturation (Robledo *et al*, 2017). NfeR1 promotes morphological changes and accelerates DNA replication, both key cell cycle-related features of bacteroid development. The precise function in this process of the cell cycle mRNAs regulated by NfeR1 remains unknown. However, genetic and transcriptomic evidence indicates that some are essential for effective symbiosis (*e.g.*, *divJ*) or are responsive to plant-derived NCR peptides that drive rhizobial differentiation into bacteroids (*e.g.*, *ftsZ* and *dnaA*) (Penterman et al, 2014; Xue and Biondi, 2019). Most notably, transcript profiling from RNA samples obtained via laser microdissection of the invasion, bacteroid differentiation, and N_2_ fixation zones of indeterminate mature alfalfa nodules revealed an overall inverse correlation between NfeR1 abundance and that of its cell cycle target mRNAs (Roux *et al*, 2014). Our data thus support NfeR1 as a post-transcriptional regulatory switch for *S. meliloti* cell cycle rewiring during infection and bacteroid differentiation, a role that merits further investigation.

### The NtrBC-NfeR1-SmelC549 regulatory axis fine-tunes *S. meliloti nod* gene expression

Nod factors, synthesized by the products of rhizobial *nod* genes in response to plant flavonoids, drive legume nodulation under N stress conditions (Honma and Ausubel, 1987; Ampe et al, 2003; Dusha and Kondorosi, 1993). The NtrBC two-component system, the master regulator of the NSR, has been reported to influence *nod* gene expression in several rhizobial species, likely through largely indirect mechanisms that remain unclear. For example, in *S. meliloti*, NtrC promotes transcription of *nodD3*, which encodes one of the three luteolin-responsive activators of *nod* gene expression (Dusha and Kondorosi, 1993). In *Rhizobium etli*, however, *ntrC* knockout reverses the negative effect of ectopic overexpression of heterologous *gdhA* on bean nodulation (Mendoza *et al*, 1995). Given its role in modulating NtrBC output and repressing *gdhA*, we reasoned that NfeR1 may also contribute to maintaining wild-type levels of *nod* gene products. Although *nod* mRNAs were not identified as direct interaction partners of NfeR1, RT-qPCR data indicate that they are positively regulated by NfeR1, primarily through *gdhA* silencing. Importantly, *gdhA* regulation by NfeR1 occurs by base pairing within the mRNA CDS and appears to be independent of NtrBC, suggesting a parallel regulatory pathway. It is likely that the accumulation of internal N, resulting from the simultaneous activity of two N assimilation pathways (GS/GOGAT and GDH), interferes with the sensing of N stress and consequently impairs *nod* gene expression. By silencing *gdhA*, NfeR1 allows the major GS/GOGAT pathway, under strict NtrBC control, to prevail, thereby ensuring proper nodulation.

A striking feature of the NfeR1 RNA interactome is the enrichment in regulatory sRNAs, all of unknown function, which significantly expands the structural and functional complexity of the NfeR1 RNA network. Among these, SmelC549, originally annotated as a *trans*-acting sRNA, was recently reclassified as a dual-function sRNA based on ribosome profiling (Ribo-seq) data, as it internally encodes the small peptide SEPr6 (Schlüter *et al*, 2010; Hadjeras *et al*, 2023). Despite the exponential increase in the identification of dual-function sRNAs in bacteria through Ribo-seq, functional characterization remains limited, with no examples yet associated with plant-microbe interactions. Our data reveal that SmelC549 is transcribed as a precursor sRNA, which is likely processed in an NfeR1-dependent manner into a readily detectable shorter isoform that accumulates under N stress conditions. The 3′ region of this abundant RNA species is largely antisense to *nodD2* mRNA and promotes accumulation of the NodD2 protein, the second of the three NodD homologs encoded in the *S. meliloti* genome (Honma and Ausubel, 1987). Notably, diversity in *nodD* copy number among rhizobial genomes has been correlated with narrower plant host range (Ferguson *et al*, 2020). Interestingly, the non-coding 3′ region of SmelC549 appears scarcely conserved across rhizobia, which may reflect target-specific evolutionary adaptations. Together, these findings suggest that NfeR1 and SmelC549 form a hierarchical regulatory module that fine-tunes *nod* gene expression, potentially contributing to host range specificity in *S. meliloti*.

Remarkably, we also detected the predicted peptide product of SmelC549, SEPr6, which is downregulated by NfeR1 through canonical base pairing at its TIR. This regulation likely occurs endogenously under N stress, when both NfeR1 and the processed regulatory form of SmelC549 reach peak levels. The inverse correlation between SmelC549 and SEPr6 abundance suggests that the regulatory and coding functions of this sRNA operate under distinct environmental contexts, N stress and mild nutrient-rich conditions, respectively. Of note, the SEPr6 coding sequence is highly conserved among rhizobial species and *A. tumefaciens*, but not in other α-proteobacteria such as *Brucella* spp., pointing to a potential role for this small peptide in plant-microbe interactions.

Our data uncover a third, unexpected role for SmelC549 as a competing endogenous sRNA (ceRNA) or RNA sponge that likely promotes decay of NfeR1 under nutrient-rich conditions (Bossi and Figueroa-Bossi, 2016; Denham, 2020). By downregulating NfeR1, SmelC549 may relieve repression on its own internal mRNA, thereby facilitating SEPr6 peptide synthesis. In parallel, SmelC549-mediated depletion of NfeR1 may prevent silencing of numerous mRNAs required to optimize *S. meliloti* fitness in mild, nutrient-rich environments. This novel NfeR1-SmelC549 RNA feedback loop provides a mechanistic explanation for how NfeR1-mediated repression of SEPr6 translation reconciles with the stability and activity of the SmelC549 regulatory form. The conservation of the NfeR1-SmelC549 interaction across plant-associated α-proteobacteria strongly suggests that this RNA-based regulatory motif is a widespread feature in rhizobia, contributing to both free-living fitness and symbiotic performance. Our findings thus provide a robust foundation for future investigations into the mechanistic and functional versatility of RNA in shaping rhizobial symbiotic behavior.

## Methods and protocols

### Bacterial strains and growth conditions

Bacterial strains and plasmids used in this work, along with their relevant characteristics, are listed in Table S2. *E. coli* strains were routinely grown in Lysogeny Broth (LB) medium at 37°C, and rhizobia in either complex tryptone-yeast (TY) or defined mannitol/glutamate minimal medium (MM) media at 30°C (Sambrook *et al*, 1989; Beringer, 1974; Robertsen *et al*, 1981). Alternatively, L-glutamate (5.4 mM) of the standard MM was replaced by 0.5 mM NH_4_Cl (MM-NH_4_). LB/MC (LB medium supplemented with 2.5 mM MgSO_4_ and 2.5 mM CaCl_2_) and MOPS-GS (50 mM MOPS, 1 mM MgSO_4_, 0.25 mM CaCl_2_, 19 mM glutamic acid, and 0.004 mM biotin, pH 7.4) media were used for cell synchronization of *S. meliloti* (De Nisco *et al*, 2014). When required, growth media were supplemented with the appropriate antibiotic(s) (µg/ml): streptomycin (Sm) 480, tetracycline (Tc) 10, and kanamycin (Km) 50 for *E. coli* and 180 for *S. meliloti*.

### DNA oligonucleotides

Sequences of the oligonucleotides used as probes for Northern hybridization, or as amplification primers for cloning and RT-qPCR are listed in Table S3.

### RNA isolation and Northern blot analysis

Total RNA was isolated from free-living bacteria cultured in the conditions described by acid phenol/chloroform extraction (Cabanes *et al*, 2000). For Northern analysis, RNA samples (15-30 µg) were subjected to electrophoresis on 6% polyacrylamide/7 M urea gels in TBE (Tris-Borate-EDTA) at ∼30 mA and electro-transferred to nylon membranes, which were subsequently probed with 5’-end radiolabeled PbNfeR1, PbSEPr6, Pb3UTRC549 or Pb5S oligonucleotides as described (del Val *et al*, 2007).

### Construction of *S. meliloti* mutants

Knockout mutants were generated by deletion of the wild-type loci using the suicide plasmid pK18*mobsacB* to induce allelic replacement by double crossover (Schafer *et al*, 1994; Torres-Quesada *et al*, 2010). Plasmids were mobilized to the parent strains by biparental mattings (Simon *et al*, 1983). SmΔ*C549* was generated in Sm2B3001 by a markerless in-frame deletion of the SmelC549 locus using pK18Δ*C549*. To construct pK18Δ*C549,* 800-bp and 775-bp DNA fragments flanking SmelC549 were amplified from genomic DNA with the EcoRIupC549/BamHIupC549 and BamHIdownC549/XbaIdownC549 primer pairs. PCR fragments were digested with *Eco*RI/*Bam*HI and *Bam*HI/*Xba*I, respectively, and ligated to the pK18*mobsacB Eco*RI and *Xba*I restriction sites, leading to insertion of the tandem fragments via their common *Bam*HI site. Similarly, SmΔ*nfeR1*Δ*gdhA* strain was generated in SmΔ*nfeR1* by a markerless in-frame deletion of the *gdhA* CDS using pK18Δ*gdhA,* including 764-bp and 788-bp DNA fragments flanking the *gdhA* ORF. For construction of pK18Δ*gdhA,* EcoRIupgdhA/XbaIupgdhA and XbaIdowngdhA/HindIIIdowngdhA primer pairs were used for DNA amplification. The PCR products were digested with *Eco*RI/*Bam*HI and *Bam*HI/*Xba*I, respectively, and then ligated to the pK18*mobsacB Eco*RI and *Xba*I restriction sites.

All PCR amplifications required for cloning were performed with the proofreading Phusion™ High-Fidelity DNA polymerase (Thermo Scientific). Plasmid inserts were always checked by sequencing to confirm the absence of PCR-introduced mutations.

### Construction of plasmids for NfeR1 tagging and induced expression

For the IPTG-induced expression of MS2 aptamer-tagged NfeR1, we constructed plasmid pSKiMS2NfeR1, which is based on the indirect *sinR-sinI* system as described previously (Robledo *et al*, 2015). NfeR1 was amplified from pSRKMS2NfeR1 (constitutively expressing MS2NfeR1) using the PCR1/PCR2 primers. The PCR product was restricted with *Xba*I and *Xho*I and inserted into pSKiMS2 to generate pSKiMS2NfeR1, which was mobilized to *S. meliloti* strain Sm2020 by biparental matings.

Replacements of specific nucleotides within the *Xba*I site were performed using a two-step PCR strategy based on overlapping fragments with pSKiMS2NfeR1 as template. The first round of PCR amplifications was performed with sinR_NdeIF or secSRK (both hybridizing to the plasmid template) and their respective primer pairs carrying the desired mutations (Mut1Fw, Mut1Rv, Mut3Fw and Mut3Rv). Each pair of complementary PCR products was used as the template in the second PCR with sinR_NdeIF/secSRK. The resulting products were digested with *Nde*I/*Xba*I and ligated to pSRKKm yielding plasmids pSKiMS2NfeR1_1 and pSKiMS2NfeR1_3.

### MAPS assays

Sm2020 cells carrying pSKiMS2NfeR1 or pSKiNfeR1 (control of column-binding specificity) were grown in MM and MM-NH_4_ media to exponential phase. Bacterial lysates were subjected to affinity purification following a previously described protocol adapted to *S. meliloti* (Robledo *et al*, 2021; García-Tomsig *et al*, 2022; Lalaouna *et al*, 2017). Briefly, cells equivalent to 200 OD_600_ were harvested by centrifugation (4°C) at 3,500 x *g* 15 min after the addition of IPTG (1 mM) to induce sRNA transcription. Cells were washed with 20 ml TE buffer (pH 8), centrifuged again, resuspended in 8 ml lysis buffer (20 mM Tris-HCl pH 8.0, 150 mM KCl, 1 mM MgCl_2_, 1 mM DTT), and broken using a French press. Soluble cell fractions were subjected to affinity chromatography on MS2-MBP-conjugated amylose resin as described (Robledo *et al*, 2021; García-Tomsig *et al*, 2022; Lalaouna *et al*, 2017). Eluted RNA was isolated by phenol/chloroform extraction followed by precipitation of the aqueous phase with 4 vol EtOH in the presence of 20 µg of glycogen. To monitor the procedure, RNA was extracted from 50 µl of cleared lysate, flow through, and wash fractions by acid phenol/chloroform extraction (Cabanes *et al*, 2000). RNA preparations were probed with PbNfeR1 upon Northern blotting.

Strand-specific cDNA libraries from RNA fractions eluted from columns were generated and sequenced in the Illumina NextSeq 500 platform. Demultiplexed sequencing reads were mapped with Bowtie2 v2.2.3 using standard parameters to the *S. meliloti* Sm1021 reference sequence downloaded from the Rhizogate portal (Becker *et al*, 2009; Langmead and Salzberg, 2012). Uniquely mapped reads were assigned to protein coding genes or non-coding RNAs with Rsubread 3.12 (Liao *et al*, 2019). Read counts for each genomic feature were normalized by coverage and the resulting RPKM values were the basis for fold change calculations (Robinson and Oshlack, 2010). The Integrative Genomics Viewer (IGV) software was used for data visualization.

### Construction of plasmids for induced SmelC549 expression

For IPTG-induced expression of wild-type SmelC549, we constructed plasmid pSKiC549, which is also based on the indirect *sinR-sinI* system (Robledo *et al*, 2015). The region encompassing *sinR-P_sinI_-TSS_sinI_*and the 400-nt long sequence from the TSS of SmelC549 were PCR amplified using genomic DNA as the template with sinR_NfeIF/TSS3_28bp_b_sinIR and C549OexfusTSSI/C549term2XbaI primer pairs, respectively. These two fragments overlap and were jointly used as template for amplification with sinR_NdeIF/secSRK. The resulting PCR product was restricted with *Nde*I and *Xba*I and inserted into pSRKKm to yield pSKiC549.

### Protein tagging with 3xFLAG

The tagging of proteins at their C-termini with three consecutive units of the FLAG epitope (3xFLAG) was achieved by using the pSK_FLAG and pR_FLAG vectors. To generate pSK*gdhA^FLAG^*, the GdhA CDS (without the stop codon) was amplified with primers NdeIgdhA/gdhAXbaI for subsequent digestion with *Nde*I/*Xba*I and cloning into pSK_FLAG.

For generation of pR*SEPr6^FLAG^* and pR*nodD2^FLAG^*, the SEPr6 and NodD2 CDSs (excluding the stop codon) were amplified using BamHITSS_C549F/ORF_C549SphIR and NheInodD2Fw/nodD2SphIRv primer pairs, respectively. These PCR products were digested with *Bam*HI/*Sph*I or *Nhe*I/*Sph*I and cloned into pR_FLAG, downstream the P*_syn_* constitutive promoter. For pR*gdhA^FLAG^* construction, the GdhA CDS and a 428-nt upstream region containing P*_gdhA_* was amplified with primers HindIIIPgdhA/gdhASphI. The DNA fragment was digested with *Hin*dIII/*Sph*I and cloned into pR_FLAG.

### Protein immunoblot

For Western blot, aliquots equivalent to 0.2 OD_600_ of cells were denatured by heating at 95°C for 5 min, resolved in 10% SDS-PAGE and blotted onto a polyvinylidene difluoride membrane (P 0.45, Amersham). Membranes were probed with a monoclonal anti-FLAG antibody (Sigma F7425; 1:5,000) as reported (Robledo *et al*, 2021). Blots were developed by incubation for 5 min in blotting detection reagent (ECL, Amersham) and signals were detected with a ChemiDoc system (BioRad). The intensity of lanes was quantified using ImageJ software (Schneider *et al*, 2012).

### Fluorescence reporter assays

The different transcriptional fusions of the NfeR1 promoter to *gfp* were individually transferred by biparental conjugation to Sm2B3001 and SmΔ*C549*. Three transconjugants of each strain were grown to exponential phase (OD_600_ of 0.2 to 0.3), divided into untreated and treated (1 mM IPTG) cultures, and incubated for 24 h and 48 h. OD_600_ and fluorescence (excitation 488 nm, emission 522 nm) were measured in a Thermo Scientific™ Varioskan™ LUX multimode microplate reader. Fluorescence values were normalized to the culture OD_600_ after subtraction of the background values from TY or MM broths.

### RT-qPCR

RNA samples obtained as described previously were additionally treated with Invitrogen TURBO^TM^ DNase for 1 h at 37°C, and further cleaned up with the RNeasy Mini Kit (Qiagen) following the manufacturer guidelines. To improve retention of the sRNA fraction, RNA samples are loaded onto the columns mixed with 7 volumes of 100% ethanol. cDNA was synthesized with the Takara Prime Script RT Master Mix (Perfect Real Time) using 1 μg of total RNA. RT-qPCR was carried out in a QuantStudio 3 (Thermo Fisher Scientific) with the Takara TB Green Premix ExTaqII (Tli RNaseH Plus) using 0.5 μl of cDNA following manufacturer’s instructions. The ratios of transcript abundance were calculated as the ΔΔCT mean average of three replicates on three independent RNA extracts, where the ΔΔCT represents the level of gene expression in the mutant strain relative to wild-type strain or in the IPTG-induced strain relative to untreated control strain. The seemingly constitutive gene *SMc01852*, encoding a phosphofructokinase, was used to normalize gene expression (Krol and Becker, 2004). Control reactions without reverse transcriptase (–RT) in the RNA samples were simultaneously performed to confirm absence of DNA contamination.

### Cell synchronization and flow cytometry

Bacterial cell synchronization was done as described previosly (De Nisco *et al*, 2014). Three independent colonies of each *S. meliloti* strain/transconjugant were cultured in LB/MC broth until OD_600_ 0.1, washed twice with sterilized 0.85% NaCl solution and resuspended in MOPS-GC for 4.5 h. Then, bacteria were collected and resuspended in MM or TY.

At different time points, aliquots equivalent to 0.6 OD_600_ units of cells were washed with phosphate-buffered saline solution (PBS) and fixed in 4% paraformaldehyde (PFA), followed by two additional washes with PBS. For DNA content measurement, 20 μl of bacterial suspensions, previously stained with 5 μg/ml 4′,6-diamidino-2-phenylindole (DAPI), were loaded into a MACSQuant VYB flow cytometer. Instrument settings were as follows: forward scatter (FSC) photomultiplier tube (PMT) voltage, 650V; side scatter (SSC) PMT, 450V; and UV fluorescence detection channel (for DAPI), 620V. MACSQuantify^TM^ software version 3.0 was used for data acquisition and analysis.

### Electron microscopy

Aliquots equivalent to 0.3 OD_600_ units of cells were washed twice with filtered PBS. Thirty microliters of each bacterial suspension were applied to formvar-coated nickel grids (FF200-Ni-50, Electron Microscopy Sciences). Then, the grids were washed twice with filtered distilled water for 1 min, stained with 2% uranyl acetate for 3 min and air-dried for 1 h. The samples were visualized using a JEOL JEM-1011 transmission electron microscope.

### Computational methods

Venn diagrams were generated with the Venny 2.0 tool (https://bioinfogp.cnb.csic.es/tools/venny/index2.0.2.html). IntaRNA (v 3.2.0; http://rna.informatik.uni-freiburg.de:8080/) was used to predict RNA-RNA base-pairing interactions (Wright *et al*, 2014). *SmelC549* sequence alignments were generated with ClustalW implemented in BioEdit (http://www.mbio.ncsu.edu/BioEdit/bioedit.html) (Larkin *et al*, 2007). ORF Finder was employed to predict protein sequences from DNA sequences (https://www.bioinformatics.org/sms2/orf_find.html).

## Data availability

Raw sequence data that support the findings of this study have been deposited in the Sequence Read Archive (SRA) with the primary accession code PRJNA1338694.

## Acknowledgements

Alicia Barroso (Genomics Unit of Instituto de Parasitología y Biomedicina “López-Neyra”, CSIC, Granada) is acknowledged for RNAseq. Dr. Andreina Peralta-Leal and Alicia Rodríguez-Sanchez (Confocal and Transmission Electron Microscopy Service - CTEM, “Estación Experimental del Zaidín”, CSIC, Granada) are acknowledged for transmission electron microscopy images. This work was supported by grants PID2020-114782GB-I00 and PID2023-147300NB-100 funded by MCIN/AEI/10.13039/501100011033, both awarded to J.I.J.-Z. M.R. was supported by a Ramón y Cajal fellowship (RYC2022-035122-I) from MCIN/AEI. N.I.G.-T was also supported by a grant for short stay of researchers, funded by Universidad de Cantabria. N.I.G.-T and S.K.G.-G were funded by FPU fellowships FPU16/01275 and FPU21/05195, respectively, from Ministerio de Universidades. The funders had no role in study design, data collection and interpretation, or the decision to submit the work for publication.

## Author contributions

**Natalia I. García-Tomsig:** planned and performed most of the experiments, analyzed the data, and wrote the manuscript. **Sabina K. Guedes-García:** contributed to experimental discussions, performed RT-qPCR assays, and critically reviewed the manuscript. **Marta Robledo:** assisted with flow cytometry experiments, contributed to data interpretation, and critically reviewed the manuscript. **José I. Jiménez-Zurdo:** acquired funding, conceived and designed the study, supported experimental work, analyzed data, and wrote the manuscript.

---

**Figure S1.**
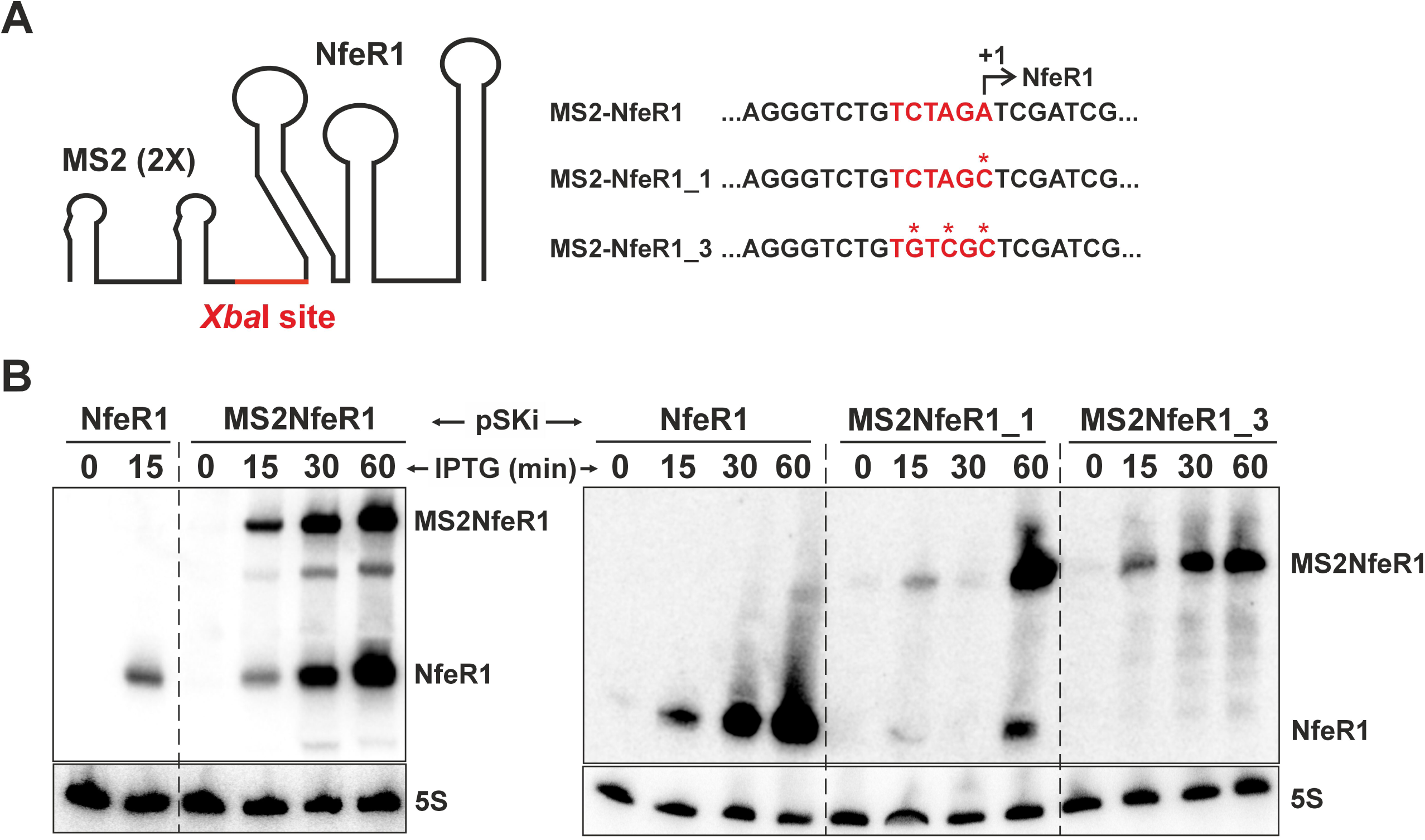
Construction of pSKiMS2NfeR1 for overexpression of MS2-tagged NfeR1. **(A)** Fusion of the MS2 aptamer to the 5’-end of NfeR1 via the *Xba*I restriction site (marked in red). The nucleotide sequence of the fusion is showed to the right. Asterisks indicate nucleotides substitutions introduced to prevent the generation of wild-type NfeR1. **(B)** Northern blot analysis of MS2NfeR1 expression. Total RNA was extracted from Sm2020 cultures in MM at 0, 15, 30 or 60 min following IPTG induction. Left panel: probing of RNA from bacteria transformed with pSKiNfeR1 or the initial construct pSKiMS2NfeR1. Right panel: probing of RNA from bacteria conjugated with pSKiNfeR1, pSKiMS2NfeR1_1 or pSKiMS2NfeR1_3. The 5S rRNA was probed as an RNA loading control.

**Figure S2.**
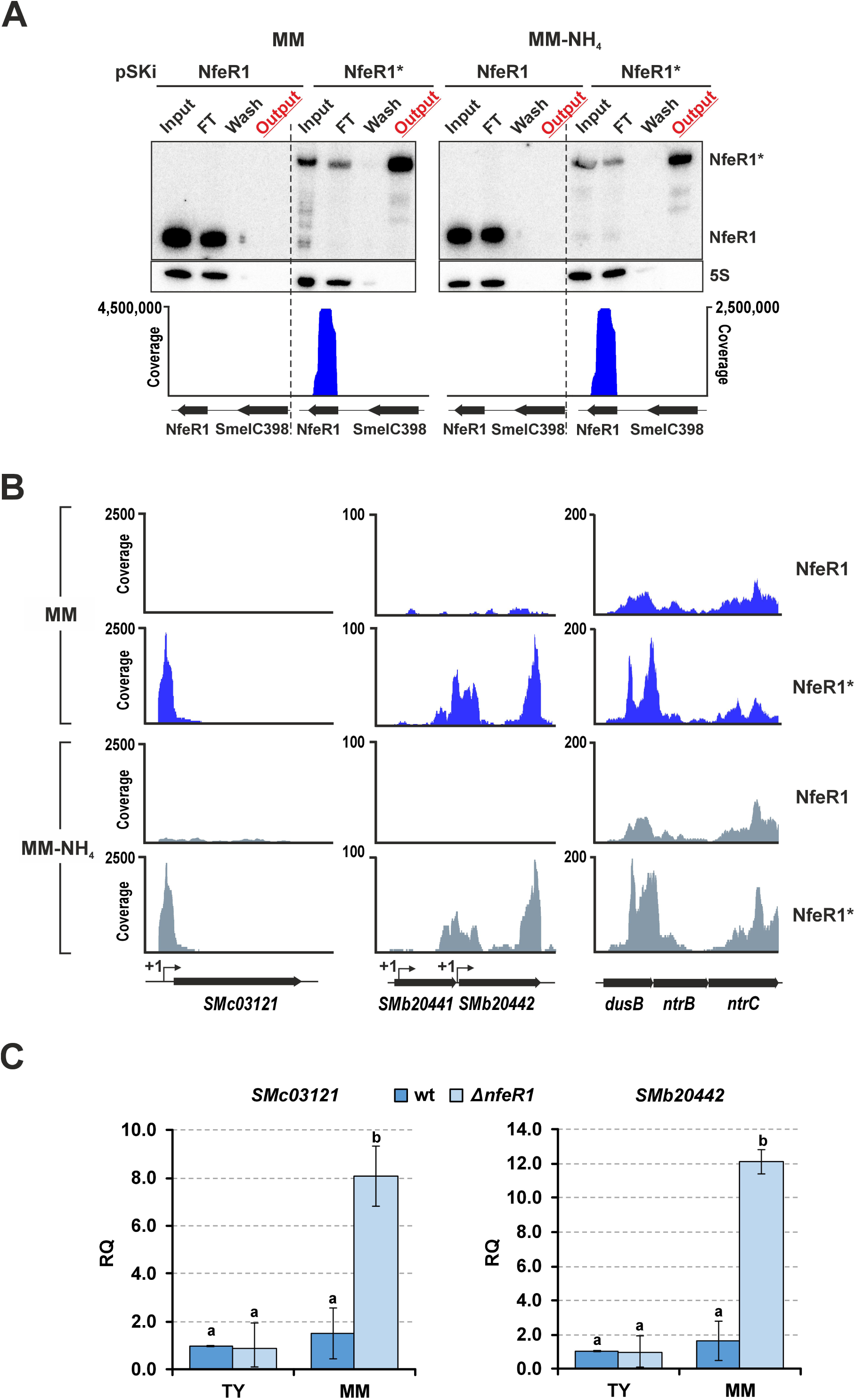
NfeR1 MAPS setup. **(A)** Monitoring of affinity chromatography in MM and MM-NH_4_. Expression of wild-type NfeR1 and MS2-tagged NfeR1 (NfeR1*) was induced for 15 min with IPTG in Sm2020 transformed with pSKiNfeR1 or pSKiMS2NfeR1, respectively. RNA from input, flowthrough (FT), wash, and output chromatography fractions (as indicated above each panel) was probed with PbNfeR1. 5S rRNA was used as a loading control. IGV plots below show of read coverage from the output fractions mapped to the NfeR1 locus. **(B)** Known NfeR1 target mRNAs co-purified efficiently with NfeR1* in MM (blue) and MM-NH_4_ (grey). IGV plots show reads coverage and recovery profiles of *SMc03121*, *SMb20442*, and *ntrBC* mRNAs following affinity chromatography using wild-type and tagged NfeR1 as baits. Known transcription start sites (TSSs) are indicated with arrows (+1). **(C)** RT-qPCR analysis of *SMc03121* and *SMb20442* mRNA abundance in the wild-type Sm2B3001 and the SmΔ*nfeR1* mutant. RNA was extracted from bacteria grown in TY to exponential phase, washed in PBS and cultured in MM for 4 h. Relative Quantification (RQ) values were normalized to *SMc01852* as a constitutive control. RQ values from MM cultures were then related to the values obtained for the wild-type strain cultured in TY. Bar graphs values are mean ± standard error of three replicates from two independent cultures (n = 6). Different letters above bars indicate statistically significant differences between groups (one-way ANOVA, *p* < 0.05, followed by post hoc Tukey’s test).

**Figure S3.**
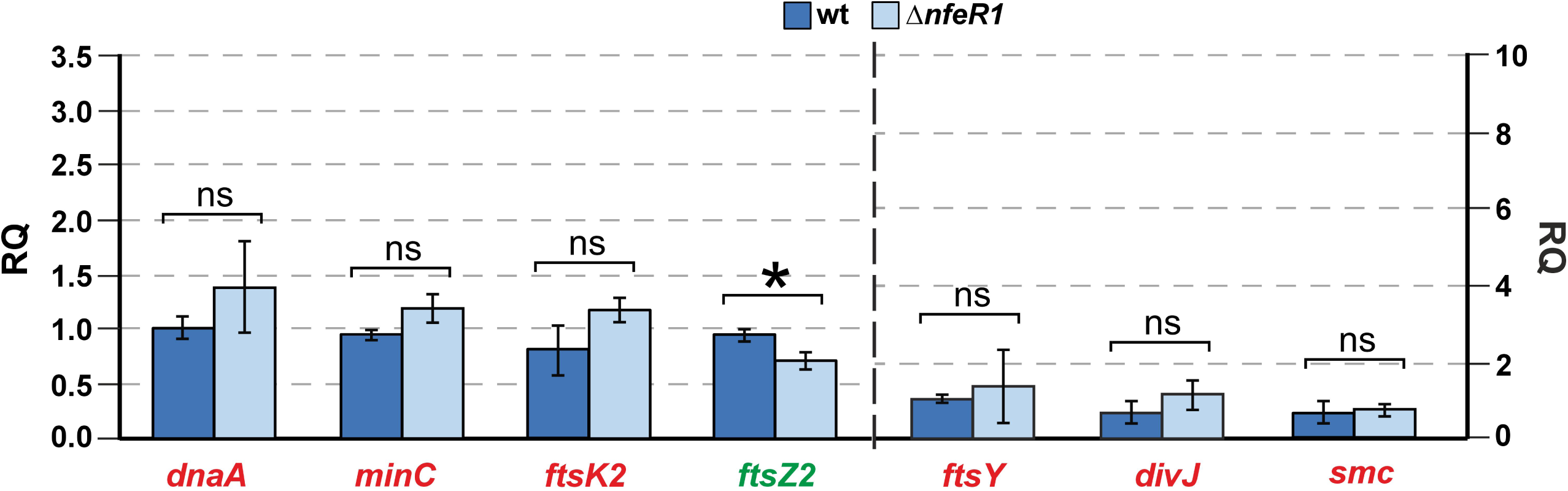
NfeR1 does not influence accumulation of cell cycle mRNAs in rich TY medium. RT-qPCR analysis of *dnaA*, *minC*, *ftsK2*, *ftsZ2*, *ftsY*, *divJ* and *smc* mRNA abundance in the wild-type Sm2B3001 strain and the SmΔ*nfeR1* mutant. RNA was extracted from bacteria grown in TY to exponential phase. Relative Quantification (RQ) values were normalized to *SMc01852* as a constitutive control. Bar graph values are mean ± standard error from three replicates of two independent cultures (n = 6). Statistical significance differences were assessed with ANOVA: ns, not significant; *, *p* < 0.01.

**Figure S4.**
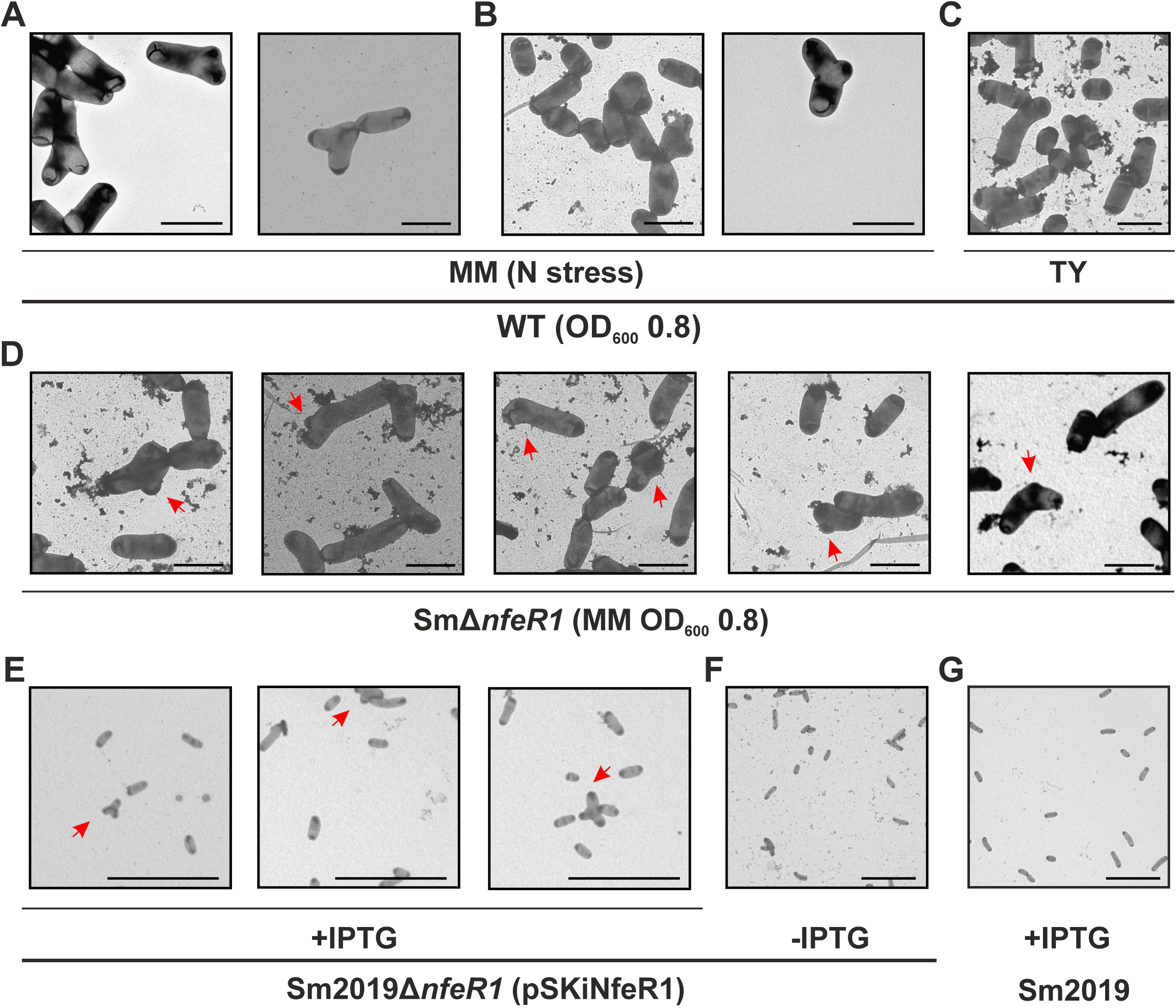
NfeR1 influences *S. meliloti* cell morphology. Different *S. meliloti* strains were cultured under nitrogen stress (MM) or non-stress (TY) conditions and examined by transmission electron microscopy (TEM) to assess changes in cell morphology. **(A)** Y–shaped morphology of wild–type bacteria grown in MM. **(B)** Less frequent morphological variants observed in MM wild–type cultures. **(C)** Canonical coccoid morphology of wild-type cells grown in TY. **(D)** Altered morphology of SmΔ*nfeR1* cells compared to the wild-type in MM. **(E)** Morphological alterations of Sm2019Δ*nfeR1* carrying plasmid pSKiNfeR1 after 16 h of IPTG-induced NfeR1 expression in TY. **(F)** Uninduced control (-IPTG) showing wild-type morphology in TY. **(G)** Wild–type strain grown in the presence of IPTG, confirming that morphological alterations observed in panel e are not due to IPTG itself. Scale bars: 2 μm (a-d) or 10 μm (e-g).

**Figure S5.**
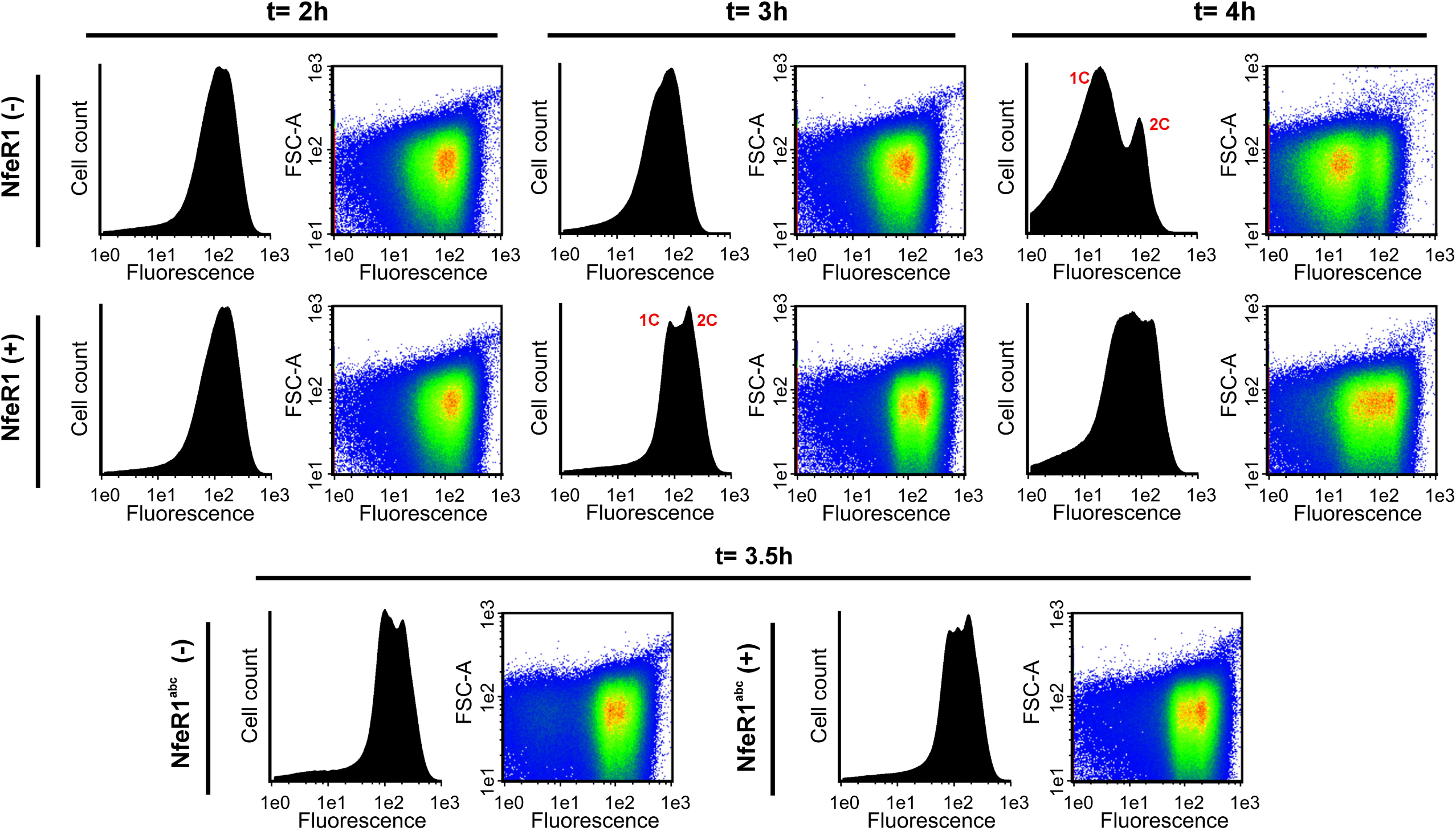
NfeR1 overexpression affects *S. meliloti* cell cycle progression. DNA content of Sm2019Δ*nfeR1* was measured by flow cytometry following IPTG-induced expression of NfeR1 or its mutant variant NfeR1abc. Synchronized cultures were resuspended in TY medium with or without IPTG. Cells were collected at various time points, stained with DAPI, and analyzed for DNA content based on fluorescence intensity. Left panels: histograms showing DNA content (X-axis) versus total cell count (Y-axis). Peaks corresponding to 1C and 2C indicate cells with one or two genome equivalents, respectively. Right panels: density plots showing the relationship between DNA content and forward scatter (FSC).

**Figure S6.**
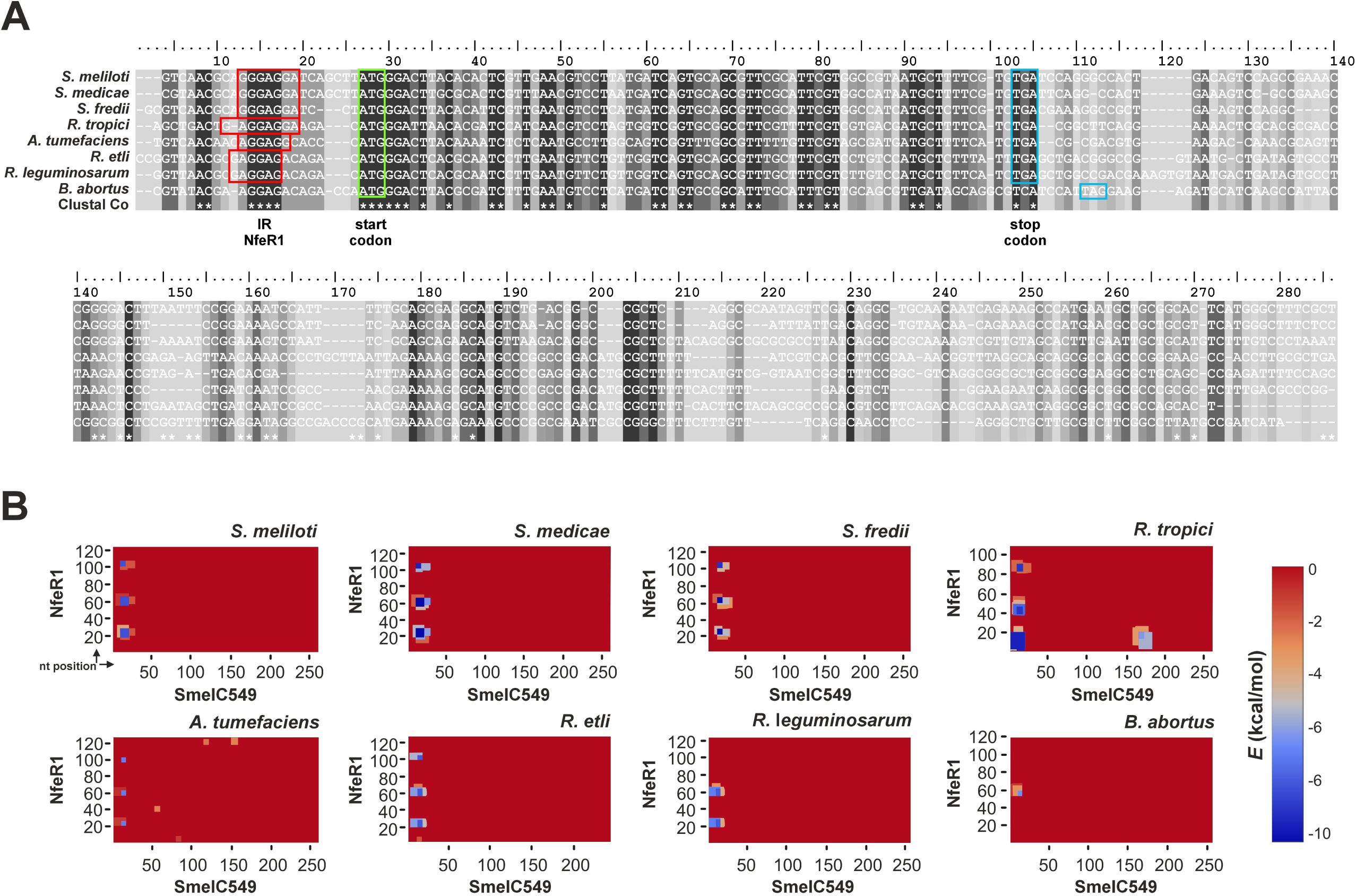
Multiple sequence alignment of SmelC549 homologs in α-proteobacteria. **(A)** Conserved sequence motifs are indicated below the alignments. The following strains encode SmelC549 homologs: *S. meliloti* 1021; *S. medicae* WSM419; *S. fredii* NGR234; *Rhizobium tropici* CIAT899; *Agrobacterium tumefaciens* C58; *R. etli* CFN42; *R. leguminosarum* bv. *viciae* 3841; *Brucella abortus* 2308. Red boxes highlight regions of putative interactions with their respective NfeR1 homologs (IR NfeR1). Green and blue boxes mark the start and stop codons of the *SEPr6* coding sequence, respectively. **(B)** Hetmaps of IntaRNA predicted base-pairing interactions between NfeR1 and SmelC549 homologs. The nucleotide position in each sRNA sequence is relative to the TSS.

**Dataset S1.** Analysis of the MAPS dataset.

**Table S1.** NfeR1-specific mRNA targets relevant to symbiosis.

**Table S2.** Bacterial strains and plasmids used in this work.

**Table S3.** Oligonucleotides specifically used in this work.

**Table S3.**
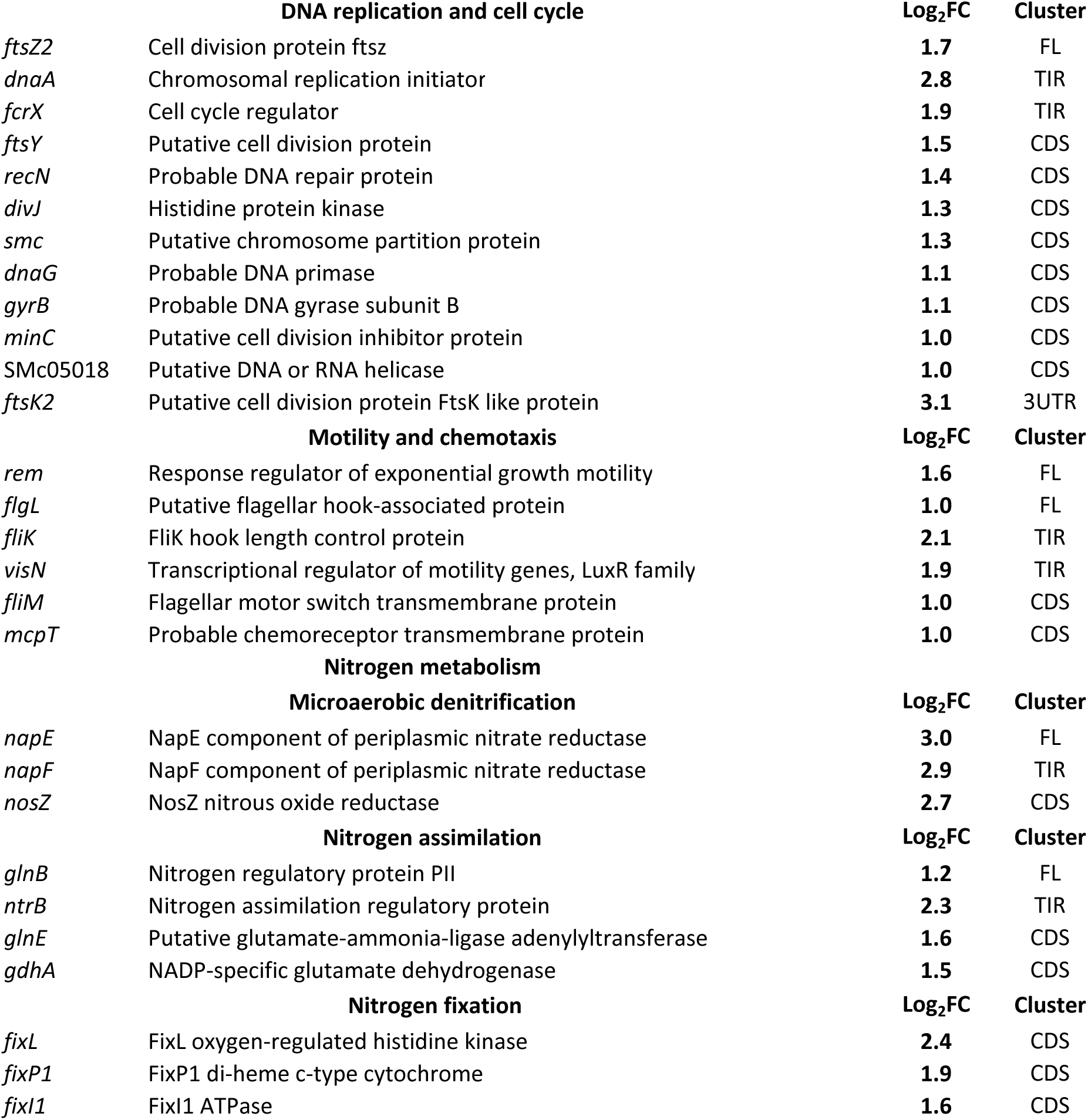
NfeR1-specific mRNA targets belonging to genes relevant to *S. meliloti* symbiosis

